# Adaptation to seasonal reproduction and temperature-associated factors drive temporal and spatial differentiation in northwest Atlantic herring despite gene flow

**DOI:** 10.1101/578484

**Authors:** Angela P. Fuentes-Pardo, Ryan Stanley, Christina Bourne, Rabindra Singh, Kim Emond, Lisa Pinkham, Jenni L. McDermid, Leif Andersson, Daniel E. Ruzzante

## Abstract

Natural selection can maintain local adaptation despite the presence of gene flow. However, the genomic basis and environmental factors involved in adaptation at fine-spatial scales are not well understood. Here, we address these questions using Atlantic herring (*Clupea harengus*), an abundant, migratory, and widely distributed marine species with substantial genomic resources including a chromosome-level genome assembly and genomic data from the eastern Atlantic and Baltic populations. We analysed whole-genome sequence and oceanographic data to examine the genetic variation of 15 spawning aggregations across the northwest Atlantic Ocean (∼1,600 km of coastline) and the association of this variation with environmental variables. We found that population structure lies in a small fraction of the genome involving adaptive genetic variants of functional importance. We discovered 10 highly differentiated genomic regions distributed across four chromosomes. Two of these loci appear to be private to the northwest, four loci share a large number of adaptive variants between northwest and northeast Atlantic, and four shared loci exhibit an outstanding diversity in haplotype composition, including a novel putative inversion on chromosome 8. Another inversion on chromosome 12 underlies a latitudinal genetic pattern discriminating populations north and south of a biogeographic transition zone on the Scotian Shelf. Our genome-environment association analysis indicates that sea water temperature during winter is the environmental factor that best correlates with the latitudinal pattern of this inversion. We conclude that the timing and geographic location of spawning and early development are under diverse selective pressures related to environmental gradients. Natural selection appears to act on early-life performance traits with differential fitness across environments. Our study highlights the role of genomic architecture, ancestral haplotypes, and selection in maintaining adaptive divergence in species with large population sizes and presumably high gene flow.

## 1. INTRODUCTION

Understanding how organisms adapt to their habitat and identifying which genes and environmental factors underpin this process are key questions in evolutionary biology and conservation. Such knowledge can help illustrate how biological diversification occurs (Nosil & Feder, 2012) and provides a genetic framework to direct management actions towards the conservation of intraspecific diversity (Hohenlohe et al., 2021). In marine species with high gene flow and large population sizes, it has been difficult to establish the extent of population structure and local adaptation by only analyzing neutral genetic markers (Hauser & Carvalho, 2008) or a small fraction of the genome. However, with the increased ability to examine both neutral and adaptive genetic variation enabled by high-throughput sequencing, recent genomic studies have revealed unprecedented levels of genetic differentiation at loci putatively under selection in marine species with seemingly undifferentiated populations at neutral loci [e.g., Atlantic herring (Lamichhaney et al., 2012), cod (Johansen et al., 2020), American lobster (Dorant et al., 2022)].

Theory proposes that gene flow can counteract adaptation by introducing maladaptive genetic variants into locally adapted populations but, at the same time, can bring in alleles that may prove beneficial in the local environment (Lenormand, 2002). This begs the question as to how local adaptation can arise and be maintained in marine species with presumably high gene flow. Empirical evidence indicates that gene flow in marine species can be restricted by a variety of ecological factors (Palumbi, 1994). These factors include differences in life history and physiological tolerance (e.g., timing of reproduction, or thermal preference/avoidance) (Lowerre-Barbieri et al., 2011), adult reproduction and behavioural phenotype favoring natal homing (Thorrold, 2001) or local recruitment (Levin, 2006), oceanography (e.g., gradients in salinity or temperature or divergent circulation patterns), and natural selection against migrants (Hendry et al., 2004; Nosil et al., 2005). When gene flow is restricted, genetic divergence can arise due to genetic drift, mutation, and selection. In marine species with large population sizes, the role of genetic drift is assumed to be minor, facilitating the detection of signatures of selection. Therefore, abundant migratory marine species inhabiting diverse environments offer ideal systems for the study of ecological adaptation with gene flow. Such is the case of Atlantic herring.

Atlantic herring is a pelagic fish that inhabits temperate waters of the north Atlantic, including fully marine environments, fjords, and the brackish waters of the Baltic Sea. It forms large schools, often comprising millions of individuals. Juveniles and adults undertake extensive annual migrations between feeding, overwintering, and spawning areas. In the northwest Atlantic (NW), spawning occurs from southern Labrador to the Gulf of Maine at relatively predictable times and locations near shore, peaking in spring (April-May) and fall (September-October) (McQuinn, 1997; Stephenson et al., 2009; Wheeler & Winters, 1984). Such diversity in reproductive phenology and spawning behaviour can promote temporal and spatial reproductive isolation between populations that are likely exposed to contrasting environments and selective pressures. These characteristics make this species an ideal model for investigating the genetic basis and mechanisms involved in ecological adaptation.

Herring plays a critical role in the marine food chains as a forage species that feeds on plankton and is an important food source for other species of fish, mammals, and sea birds. In addition, it sustains large fisheries throughout the north Atlantic (FAO, 2019), some of which have experienced periods of severe decline in the last century (Engelhard & Heino, 2004; Overholtz, 2002; Simmonds, 2007). In the NW Atlantic, declining abundance has impacts on both commercial landings and the various fisheries that use forage fish such as Atlantic herring and Atlantic mackerel (*Scomber scombrus*) as bait. This is the case of Atlantic lobster (*Homarus americanus*), which has become the most valuable fishery resource in North America. In 2022, following a lack of recovery, Fisheries and Oceans Canada (DFO) announced a moratorium on directed commercial and bait fisheries for the spring stock of Atlantic herring in the southern Gulf of St. Lawrence (DFO, 2022). Though the status of individual stocks is variable across the Atlantic Canadian region (e.g., from cautious to critical in the Gulf of St Lawrence, or stable in Newfoundland), herring abundance overall has been decreasing or sustained at low levels, which is largely attributed to high adult mortality and low recruitment (Turcotte et al., 2021). Therefore, in recent years there has been increased interest in understanding the population dynamics of herring stocks using diverse approaches, including genetics.

Numerous research efforts have aimed to estimate the level of population structure of herring using diverse genetic markers and at various spatial scales, primarily in the northeast (NE) Atlantic. Initial genetic studies analyzed a few dozens of neutral markers and reported panmixia or low population differentiation (Andersson et al., 1981; André et al., 2011; Bekkevold et al., 2005; Jørgensen et al., 2005; Ruzzante et al., 2006). Subsequent studies based on genetic variants derived from the transcriptome (Limborg et al., 2012) or reduced-representation sequencing (Guo et al., 2016) or that used genome sequencing and an exome assembly (Lamichhaney et al., 2012) showed that genetic differentiation occurs at outlier loci likely associated with environmental gradients in the Baltic. Whole-genome studies using a scaffolded reference genome revealed a number of genes associated with ecological adaptation (Martinez Barrio et al., 2016), some of which are shared between European and Canadian populations (Lamichhaney et al., 2017). The recent assembly of a chromosome-level reference genome for the species enabled the possibility to examine structural rearrangements and more complete gene models (Pettersson et al., 2019), a breakthrough in the ability to study the genetic basis of adaptation. The most recent whole-genome study analysed 53 locations, 48 from the NE and 5 from the NW Atlantic, and disclosed hundreds of loci underlying ecological adaptation (Han et al., 2020). About 30 loci showed consistent association with adaptation to salinity, seven to spawning time, and four putative chromosomal inversions are presumably related to adaptation to water temperature during spawning, which is higher in waters around southern UK and Ireland. Candidate genes associated with spawning time include the thyroid-stimulating hormone receptor (*tshr*), *sox11b* transcription factor (*sox11b*), calmodulin (*calm1b*), and estrogen receptor 2 (*esr2a*), all located on chr 15 and with a known or presumed role in reproductive biology. The gene *myhc* (myosin heavy chain) on chr 12 is putatively involved in myogenesis and plasticity of seasonal development in herring (Johnston et al., 2001).

A few initiatives have focused on the population structure of NW herring Atlantic. Some studies analysed a dozen of microsatellite markers (McPherson, Stephenson, O’Reilly, & Jones, 2001; McPherson, Stephenson, & Taggart, 2003; McPherson, O’Reilly, & Taggart, 2004). One compared samples collected at the same location 10 years apart and found temporal stability of allele frequencies at a subset of SNPs related with spawning time (Kerr et al., 2019). Others have used whole genome approaches primarily focused on NE Atlantic populations in addition to the same five Canadian populations (Han et al., 2020; Lamichhaney et al., 2017). Low genome-wide differentiation at presumed selectively neutral loci was found across studies. However, one of the whole-genome studies showed that about 25% of SNPs associated with seasonal reproduction are shared between NW and NE Atlantic populations (Lamichhaney et al., 2017). The fact that a large proportion of outlier loci are likely not shared between populations across the ocean, raises the question of whether herring display local adaptation to NW Atlantic environments.

Here, we test this hypothesis through an exhaustive examination of genome-wide variation of 15 spawning aggregations of herring in the NW Atlantic. In particular, we address three focal questions: 1) what is the temporal and spatial scale of population structuring in the focal area? 2) what is the genomic basis of population divergence and local adaptation? and 3) which evolutionary mechanisms and environmental factors are likely involved in population divergence? We integrate pooled DNA whole-genome sequencing (pool-seq) and oceanographic data to address these questions. Given the role of temperature and photoperiod in regulating the onset of seasonal reproduction in numerous temperate fish species (Migaud et al., 2010) and the relatively high predictability of spawning season and location in Atlantic herring (Sinclair & Tremblay, 1984), we expect that genetic differences among aggregations are largely due to natural selection.

## 2. MATERIALS AND METHODS

### 2.1 Study area and sample collection

The study area comprises the reproductive range of *C. harengus* in the NW Atlantic, from southern Labrador in Canada to the Gulf of Maine in the U.S. (Figure 1, Table 1). We collected individuals from 10 locations across this area. Sample collection took place between 2012 and 2016 during or near the peak of spawning at each site, mainly in spring (April to May), summer (June) and fall (August to October) seasons. We targeted individuals actively spawning or ready to spawn (gonadal maturity stages 5 and 6, respectively) when possible, to have a representative sample of distinct reproductive units (or populations) and to minimize the chance of sampling non-spawning migrants. Most of the fish were actively spawning or ready to spawn, except in the Bras D’Or lake (BDO-M), in which only 15% of fish were mature and the rest were in resting condition (Figure S1). Therefore, this sample was considered mixed. Gonadal maturity was visually assessed following classifications of Bucholtz, Tomkiewicz, & Dalskov (2008). Muscle or fin tissues were collected from each individual and preserved in 95% ethanol at −20 °C until processing.

**Figure 1.**
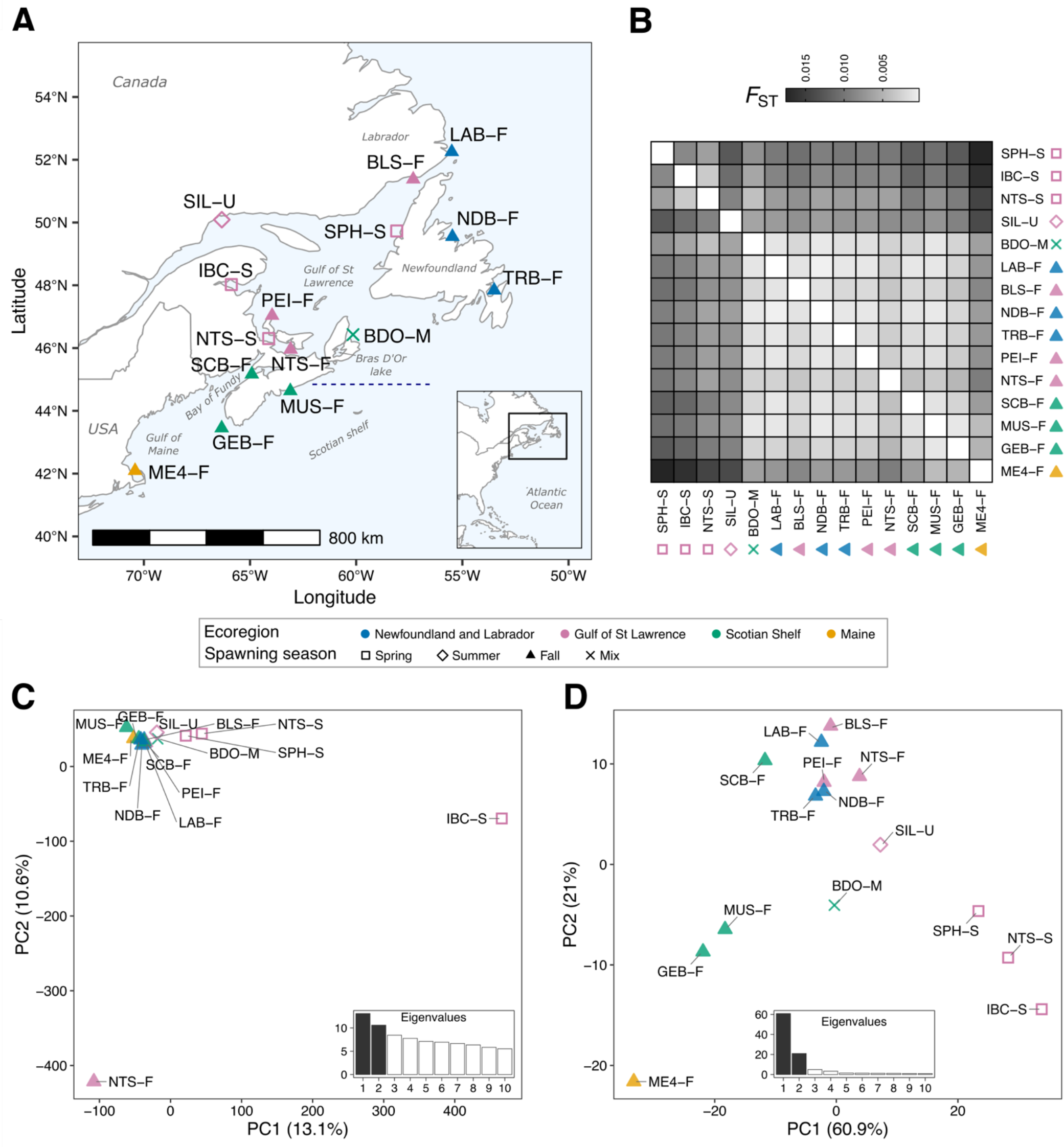
Sampling locations and population structure of northwest Atlantic herring. (**A**) Map depicting collection sites. Location name abbreviations as described in Table 1, in which the spawning season is indicated with the suffix “-S” for spring, “-F” for fall, “-U” for summer, and “-M” for mixed. Each point corresponds to a pool sample. Point colors indicate the designation of the sample to one of the major biogeographic units described for the Canadian Atlantic Ocean (DFO, 2009). Symbol shapes represent the predominant spawning season based on individual gonadal maturation status at the time of collection (Figure S1). The horizontal blue dashed line on the map indicates the approximate location of a biogeographic transition zone in the eastern Scotian Shelf, 44.61°N ± 0.25 (Stanley et al., 2018). (**B**) Heatmap plot representing pairwise *F*_ST_ estimates based on pool allele frequencies of 5,073,572 SNPs (Table S2). Samples are ordered by season, and within season, by latitude. Cell shading represents the degree of genomic divergence between a pair of pool samples, which goes from lack (white) to high (black) differentiation. Principal component analysis plot based on undifferentiated (**C**) (n = 135 950) and highly differentiated (**D**) markers (n = 545). In both plots the first two axes or principal components (PCs) are shown, and the inset bar plots indicate the percentage of the variance explained by each of the first 10 PCs. Same as before, each point corresponds to a pool sample, its color indicates the designated biogeographic unit and its shape, the corresponding spawning season.

**Table 1.** Characteristics of the 15 herring spawning aggregations included in this study.

| Locality | ID code | Sample size (N) | Longitude | Latitude | Sampling date | Spawning season | Source | BioSample accession |
| --- | --- | --- | --- | --- | --- | --- | --- | --- |
| Seven Islands (Sept-Îles) | SIL-U | 50 | -66.33 | 50.09 | 06 June 2012 | Summer | This study | SAMN33007718 |
| Inner Bay Chaleur | IBC-S | 41 | -65.85 | 48.00 | 08 May 2014 | Spring | (Lamichhane et al., 2017), WInB | SAMN05589078 |
| Stephenville | SPH-S | 48 | -57.94 | 49.73 | 30 May 2012 | Spring | This study | SAMN33007720 |
| Northumberland Strait | NTS-S | 49 | -64.12 | 46.30 | 06 May 2014 | Spring | (Lamichhane et al., 2017), WNsS | SAMN05589080 |
| Northumberland Strait | NTS-F | 50 | -62.60 | 45.73 | 16 Sep 2014 | Fall | (Lamichhane et al., 2017), WNsF | SAMN05589079 |
| Labrador | LAB-F | 50 | -55.50 | 52.25 | 24 Aug 2014, 22 Aug 2015 | Fall | This study | SAMN33007723 |
| Blanc Sablon | BLS-F | 49 | -57.31 | 51.38 | 13 Aug 2014 | Fall | This study | SAMN33007724 |
| Notre Dame Bay | NDB-F | 50 | -55.47 | 49.55 | 26 Oct 2015 | Fall | This study | SAMN33007725 |
| Trinity Bay | TRB-F | 49 | -53.47 | 47.84 | 28 Sep 2014 | Fall | (Lamichhane et al., 2017), WBob | SAMN05589075 |
| Prince Edward Island | PEI-F | 50 | -63.96 | 47.04 | 25 Aug 2014 | Fall | This study | SAMN33007727 |
| Bras D'Or lake | BDO-M | 50 | -60.85 | 45.93 | 20 April 2016 | Spring | This study | SAMN33007728 |
| Scots Bays | SCB-F | 50 | -64.92 | 45.17 | 24 Aug 2015 | Fall | This study | SAMN33007729 |
| Musquodoboit | MUS-F | 50 | -63.10 | 44.63 | 28 Oct 2015 | Fall | This study | SAMN33007730 |
| German Banks | GEB-F | 48 | -66.18 | 43.16 | 28 Aug 2014 | Fall | (Lamichhane et al., 2017), WGeB | SAMN05589077 |
| Maine fishing area 514 | ME4-F | 50 | -70.41 | 42.09 | 19 Oct 2015 | Fall | This study | SAMN33007732 |

### 2.2 DNA extraction and pool-sequencing

Total genomic DNA was isolated from individual tissues using a standard phenol chloroform protocol (Sambrook & Russell, 2006). DNA concentration (in nanograms per microliter, ng/µl) was measured in triplicates using the Quant-iT PicoGreen dsDNA assay (Thermo Fisher Scientific, U.S.) and the Roche LightCycler 480 Instrument (Roche Molecular Systems, Inc., Germany). DNA integrity was inspected using 0.8% agarose gel electrophoresis with 0.5x TBE buffer and a 1 kilo base pairs (Kbp) molecular weight ladder.

Sequence data of each spawning aggregation was obtained using the pool DNA whole genome sequencing (pool-seq) approach. This technique involves sequencing the combined DNA of several individuals from a population using a single-barcoded library to generate population-level whole genome data for a fraction of the cost of sequencing individuals to high depth, at the expense of missing individual genotypes (Schlötterer et al., 2014). We aimed to mix in a pool equal amount of DNA of 50 individuals collected at a given spawning site and season. Individual DNA were normalized to a common concentration of 30 ng/µl and combined to a single tube using the liquid handling robot epmotion 5407 (Eppendorf, Germany). Sequencing library preparation and high-throughput sequencing were performed by The Centre for Applied Genomics of the Hospital for Sick Children, Canada. A TruSeq Nano Illumina DNA library with an insert size of 550 bp was built for each DNA pool. Illumina paired-end reads (2 x 126 bp) were generated for each library with a HiSeq-2500 sequencer, targeting a 50x average depth of coverage per pool. We combined this dataset with pool-seq data from five populations that were studied before (Lamichhaney et al., 2017) (Table 1). Therefore, the northwest Atlantic was represented by a total of 15 pool samples.

### 2.3 Sequence filtering, alignment, and variant calling

Sequence quality of raw reads per pool was examined with *FastQC* v0.11.5 (Andrews, 2010), and a single report for all pools was generated with *MultiQC* v.1.3 (Ewels et al., 2016). Illumina adapters, low quality bases (Phred score < 20), reads shorter than 40 bp, and single-end reads were removed from our dataset using *Trimmomatic* v.0.36 (Bolger et al., 2014) [parameters: ILLUMINACLIP:TruSeq3-PE-2.fa:2:30:10 SLIDINGWINDOW:5:20 MINLEN:40].

The filtered paired-end sequences were mapped against the chromosome-level genome assembly of Atlantic herring (Pettersson et al., 2019) using the Burrows-Wheeler Aligner (*BWA*) v0.7.12-r1039 [default parameters, MEM algorithm] (Li, 2013). Read mapping quality was assessed with *Qualimap* v.2.2.1 (Okonechnikov et al., 2015). Mapped reads in the form of BAM files were sorted, PCR duplicates marked, and read groups added using *Picard tools* v2.10.2 (Broad Institute, 2018). An index file for each BAM file was generated with *SAMtools* v1.5 (Li & Durbin, 2009).

We called variants using the *UnifiedGenotyper* algorithm implemented in the program *GATK* v3.8 (McKenna et al., 2010). Indels were removed and biallelic SNPs were kept for further filtering. We required SNPs passed *GATK* variant quality filters, which assess sequence and mapping quality, strand bias, and variant position on reads to identify high quality polymorphisms and discard spurious ones. We established cut-offs values for the *GATK* quality filters based on density plots (Figure S2) [filters applied: FS > 60.0, SOR > 3.0, MQ < 40.0, MQRankSum < −12.5, ReadPosRankSum < −8.0] (for a description of each annotation see (Broad Institute, 2016)). Using *BCFtools* (Danecek et al., 2021), we retained SNPs that had a missing rate < 20%, a minimum minor allele count of 2, and were polymorphic. We retained the SNPs within the 5-99 percentile of the coverage distribution per pool (Figure S3), thus excluding spurious SNPs in repetitive regions and copy number variants that commonly have excessively high coverage.

Allele frequencies in pool-seq are derived from the read counts of each allele per variant site. Read counts can vary among pools and along the genome due to technical biases during sequencing (Dohm et al., 2008; Kolaczkowski et al., 2011). To account for sampling error of the pool during sequencing before calculating allele frequencies, we rescaled the raw read counts to the effective sample size *n*_eff_ using a python script that implemented the formula:

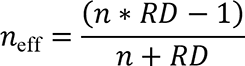

where *RD* is the raw read depth and *n* is the number of chromosomes in a pool (e.g., *n* = 2\**number of individuals in a pool*, for diploid species). *n*_eff_ represents an estimate of the effective number of chromosomes in a pool adjusted by the read depth (Bergland et al., 2014; Feder et al., 2012; Kolaczkowski et al., 2011). Based on the rescaled read counts, we calculated allele frequencies per pool.

### 2.4 Population structure and genetic diversity

We estimated the genomic differentiation between pool samples using pairwise-*F*_ST_ and a Principal Components Analysis (PCA). We computed the unbiased *F*_ST_ for pools (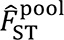) between all pairs of spawning aggregations using the *R* package *poolfstat* (Hivert, 2018). This algorithm calculates *F*-statistics equivalent to Weir & Cockerham (1984) and accounts for sampling error in pool-seq.

To compare population structure patterns of undifferentiated (presumed neutral) and highly differentiated loci (presumed under selection), we separately performed PCA on two SNP sets defined from the empirical distribution and standard deviation (SD) of allele frequencies of all markers, as applied in Han et al. (2020). The set of undifferentiated markers corresponded to SNPs with allele frequency below and close to the mean value (0.03 < allele frequency SD ≤ 0.09), while the highly differentiated markers had an allele frequency SD ≥ 0.2 from the mean (Figure S4). To reduce the effect of physical linkage among loci, we retained one SNP every 1 Kbp for the undifferentiated markers, and one SNP every 10 Kbp for the differentiated markers assuming higher linkage in regions under selection.

To characterize the genetic diversity within populations, we estimated the unbiased nucleotide diversity (π) and Tajima’s D for each pool using the program PoPoolation 1.2.2 (Kofler et al., 2011). In brief, a pileup file was generated from the BAM file of each pool using samtools v.1.10 (Li, 2011). Indels and SNPs 5 bp around indels were removed. To account for coverage variation due to sequencing, we subsampled without replacement the depth of coverage of each pileup file to a uniform value, corresponding to the 5% quantile of the coverage distribution (the minimum coverage of a SNP to be retained). SNPs with read depth between 5- 99% of the empirical coverage distribution per pool, minimum base and mapping quality of 20, and a minor allele count of 2 were retained for further analyses. The statistics π and Tajima’s D were calculated in sliding windows of 10 Kbp with a step size of 2 Kbp. Windows with a coverage fraction ≥ 0.5 were used for further analyses. All statistical tests and graphics were performed in *R* (R Core Development Team, 2023).

### 2.5 Detection of genomic regions under selection

To identify outlier genomic regions that are likely targets of selection, we calculated the absolute difference in allele frequencies per SNP (delta allele frequency, dAF) between paired contrasts of pools, using the formula: *dAF = absolute[meanAF(group1) – meanAF(group2)]*. Based on PCA clustering patterns and prior ecological knowledge of spawning stocks, the two most informative contrasts were established to compare: (i) spring and fall spawners, and (ii) northern and southern fall spawners. This analysis was complemented with the calculation of the moving average (or rolling mean) of dAF values over 100 consecutive makers, which smooths out the signal of divergence while ruling out the effect of single outlier SNPs that might result from sequencing bias.

To identify novel loci in the NW Atlantic that are likely related with local adaptation, we compared the outlier SNPs found in this study with those reported in Han et al. (2020), showing strong and consistent association with adaptation to spawning time and salinity among northeast Atlantic and Baltic populations, and within four inversions (on chr 6, 12, 17, and 23). We visually inspected the allele frequencies of outlier SNPs in our 15 NW Atlantic populations and in previously published data of one Pacific herring and 47 Atlantic herring pools (Han et al., 2020) (Table S3). In addition, we generated a neighbor-joining tree based on the allele frequencies of outlier SNPs for all 63 populations.

### 2.6 Genome-environment association and isolation-by-distance

To identify which environmental variables are strongly correlated with putatively adaptative SNPs underlying a latitudinal genetic pattern, we performed a redundancy analysis (RDA). The environmental dataset consisted of hours of day light (dayLightHours), sea surface temperature (SST) and salinity (SSS) during the months of spawning at each location, and for the winter and summer seasons, representing the coldest and warmest periods of the year. These variables were chosen because they are relevant in fish physiology and have been linked to population structuring of numerous marine species in the NW Atlantic (Stanley et al., 2018). Temperature and salinity values for each sample location were derived from monthly environmental data layers of SST and SSS developed for the North Atlantic between 2008-2017 based on the Bedford Institute of Oceanography North Atlantic Model (BNAM), a high-resolution numerical ocean model. A detailed description of oceanic (Madec et al., 1998) and sea ice (Fichefet & Maqueda, 1997) components of the model can be found in Wang, Brickman, Greenan, & Yashayaev (2016) and Brickman, Hebert, & Wang (2018). Data layers were converted to an ASCII grid with a NAD83 projection (ellipse GRS80), and had a nominal resolution of 1/12° (∼5 km^2^). Binned data for the months of spawning at each location (spring: April-May, fall: September-October), and for the winter (January-February-March) and summer (July-August-September) seasons, were averaged across 9 years in order to capture long-term trends of oceanographic variation. Extractions (mean, min, and max values) were taken from these averaged layers for each sample location. Prior to performing RDA, environmental data were standardized to zero mean and unit variance using R. Collinearity between environmental variables was assessed using two methods, with pairwise correlation coefficients computed with the function *pairs.panels* of the R package *psych* (Revelle, 2018) (Figure S5), and with variance inflation factors (VIF) obtained from RDA models built with the R package *vegan* (Dixon, 2003).

The most collinear variables were removed, and remaining collinear variables were identified and excluded one-by-one in consecutive RDA runs based on their VIF value. The variable with the highest VIF was discarded in each run until all variables had a VIF < 5, following recommendations of Zuur, Ieno, & Elphick (2010). The uncorrelated set of environmental variables consisted of summer sea surface temperature (SST_Summer_), winter sea surface temperature (SST_Winter_), sea surface temperature during spawning (SST_Spawn_), and day light hours (Figure S6).

For RDA, we used the uncorrelated set of environmental data as constraining factors for the population allele frequencies of the SNPs within the chromosomal inversion on chr 12 (chr12:17823410-25605433). RDA runs were performed with the R package *vegan*. Environmental variables that best explained the genetic variance were confirmed with a bi-directional stepwise permutational ordination method (1000 iterations) using the function *ordistep*. Significance of the overall RDA model and the most explanatory environmental variables was assessed with an analysis of variance (ANOVA) using 9999 permutations.

To evaluate whether the latitudinal genetic pattern follows an isolation-by-distance (IBD) pattern, we calculated the significance of the linear relationship between geographic and genetic distances between pairs of locations for all SNPs and for only the ones associated with the latitudinal genetic pattern (within the chr 12 inversion). For this, we performed separate Mantel tests (Mantel 1967) with 9999 permutations for each set of loci using the R package *ade4* (Dray & Dufour, 2007). Genetic distances were linearized 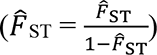 (Rousset, 1997) and 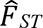 estimates were computed either using all SNPs identified across the genome, or solely the outlier SNPs strongly associated with latitudinal divergence. Geographic distances were estimated with the R package *CartDist* (Stanley & Jeffery, 2017) as the least-coast oceanic distance in kilometers (km) where land is treated as a barrier to movement.

## 3. RESULTS

### 3.1 Population-scale whole-genome sequencing

We collected 697 individuals from 10 spawning aggregations throughout the NW Atlantic (Figure 1A). We pooled DNA of 41-50 individuals per location and generated paired-end short reads for the whole genome using a HiSeq Illumina sequencer. Our sequencing effort yielded a total of ∼200 Giga base pairs (Gb) of data. The median depth of coverage per pool ranged between 57X and 77X (Table S1). We combined the pool-seq data of these 10 populations with data of five Canadian populations previously published (Han et al., 2020), for a total of 15 pools constituting the dataset of northwest Atlantic herring. After mapping reads against the genome assembly of the Atlantic herring (Pettersson et al., 2019), calling variants, and applying quality filters to the raw variants, we obtained a total of 5,264,683 high quality SNPs.

### 3.2 Population structure

We investigated the population structure among samples using pairwise-*F*_ST_ and PCA. The *F*_ST_ values of pools (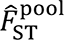) ranged between 0.001 and 0.018 (Figure 1B, Table S2), indicating low genomic differentiation among spawning aggregations across ∼1,600 km of coastline. This result is consistent with previous studies of Atlantic herring (Han et al., 2020). Despite low differentiation, we detected two subtle patterns of genetic structure between: (i) spring and fall spawners (SPH-S, IBC-S, NTS-S vs. others, mean 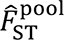 0.010 ± 0.001); (ii) the southernmost sample in the Gulf of Maine (ME4-F) and the other fall spawners (mean 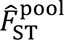 0.007 ± 0.001).

To evaluate whether population structure patterns were mainly explained by undifferentiated (putatively neutral) or highly differentiated loci (putatively selective), we separately performed PCA based on the allele frequencies of a subset of neutral (n = 135,950, Figure 1C) or outlier markers (n = 545, Figure 1D) (see Methods for details). In the PCA with neutral markers, most populations formed a single tight cluster with the exception of two samples from the Gulf of St Lawrence (IBC-S and NTS-F), which appeared as outliers. The first two principal components (PCs) explained 23.7% of the genetic variance, which are largely driven by these two samples.

In stark contrast, in the PCA based on presumably adaptive markers, spring and fall spawning populations separated on PC1, and fall spawners separated following a latitudinal pattern along PC2. These two PCs explained a large proportion of the genetic variance (82.0%). On PC2, the “northern” samples included populations from Labrador (LAB-F), Newfoundland (BLS-F, TRB-F, NDB-F), the Gulf of St Lawrence (NTS-F, PEI-F), and the Bay of Fundy (SCB-F), while the “southern” samples comprised populations from the Scotian Shelf (GEB-F, MUS-F) and the Gulf of Maine (ME4-F). On PC1, populations from Bras D’Or lake (BDO-M) and Sept-Îles (SIL-U), the first being a mixed sample and the latter was collected in the summer, appear to be closer to fall spawning samples. On PC2, spring samples were separated from each other and from the summer sample from Sept-Îles (SIL-U), in concordance with the pairwise 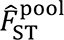 estimates.

### 3.3 Genomic regions under selection

Despite low genome-wide population structure, we uncovered genomic regions displaying significant differentiation between populations. These regions were detected through genome scans based on the absolute difference in allele frequencies (dAF) of 5,264,683 SNPs between pool samples representing distinctive ecotypes or clusters in the PCA. The informative comparisons were: (i) spring vs. fall spawners, and (ii) northern vs. southern fall spawners.

We discovered that the genomic regions showing the strongest association with spawning time were confined to nine major loci distributed on four chromosomes (chr): chr 8 (one locus), chr 12 (one locus), chr 15 (five loci), and chr 19 (two loci) (n = 2137 SNPs, Figure 2A). Four of these loci are considered “novel”, as they are not among the loci reported in Han et al. (2020) as significantly associated with ecological adaptation (chr8:23040136-30729461, chr15:6750000- 7000000, chr15:9200000-9350000, chr19:23155000-23300000). While some SNPs in these loci have been associated with spawning time before (Han et al., 2020) (n = 387, blue triangles in Figure 2A), we discovered a large number of outlier SNPs (n = 1748, 81.8%) (Figure 3, 4, red dots) that showed a stronger genetic differentiation in the west than in the east Atlantic. Most of these SNPs were located in a putative structural variant (SV) on chr 8 (n = 1349, 77% of novel SNPs) (Figure 3) that has not been reported previously. Other novel outliers SNPs were found in loci with known association with spawning time on chr 12, 15, and 19 (Figure 4).

**Figure 2.**
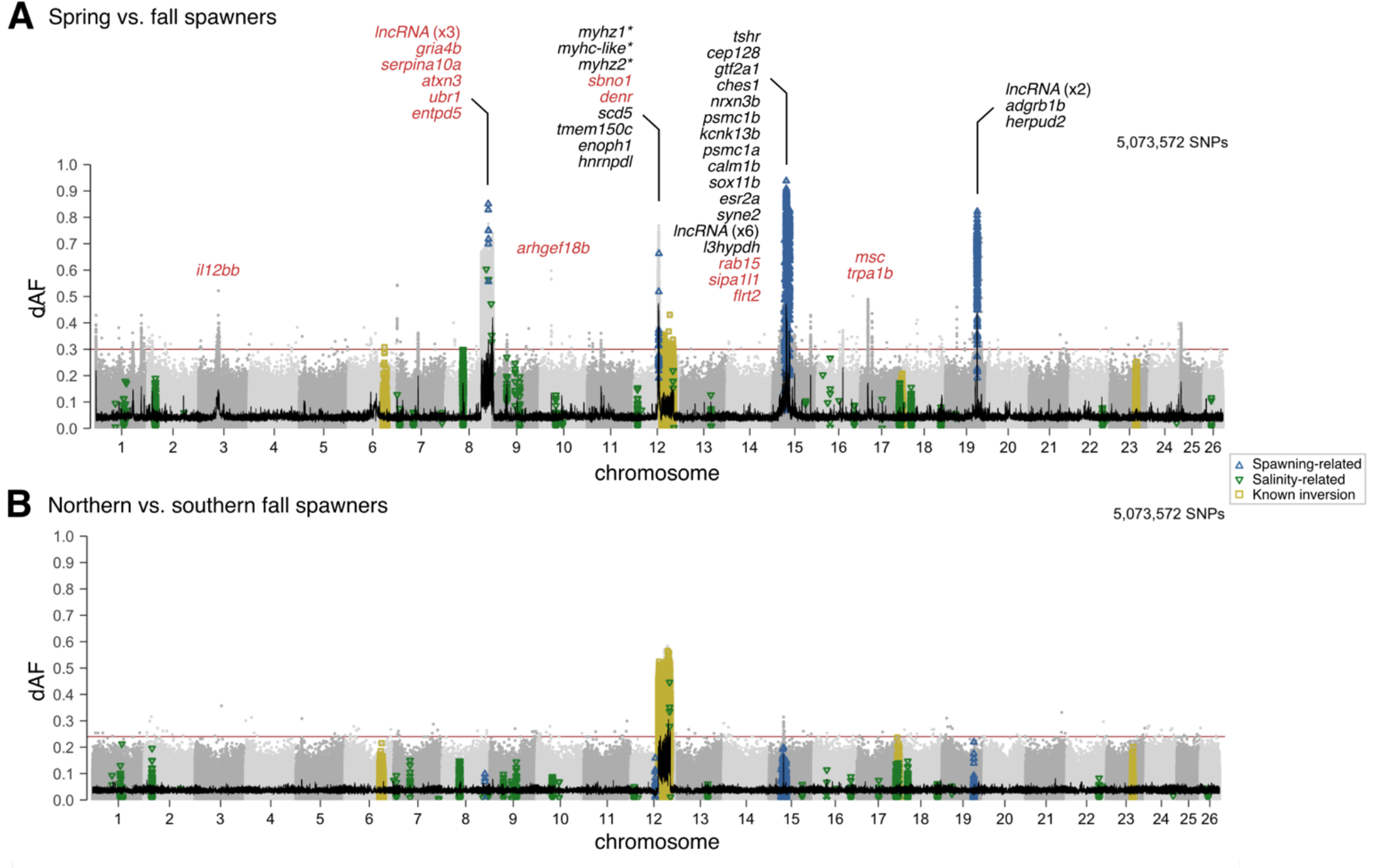
Genomic regions associated with adaptation to seasonal reproduction and a latitudinal environmental cline. Genetic differentiation (dAF) across the genome (**A**) between spring and fall spawners, and (**B**) between fall spawners in the north vs. south of the transition zone in eastern Scotian Shelf (Stanley et al., 2018). Each dot represents a single SNP. SNPs previously reported in (Han et al., 2020) as strongly associated with spawning season, salinity, and four known inversions (on chr 6, 12, 17, and 23) are denoted as empty blue upward triangles, green downward triangles, and yellow squares, respectively. Consecutive chromosomes are colored in intercalating shades of gray. The 100 SNP-rolling average of dAF is shown as a black line and the Bonferroni significance value across the genome is shown as a horizontal red line. Genes within ± 40 Kbp of the most divergent SNPs are shown on top of each informative locus. Gene names with an were inferred from homology with orthologous genes. Gene names colored in red correspond to genes within the associated genomic region.

**Figure 3.**
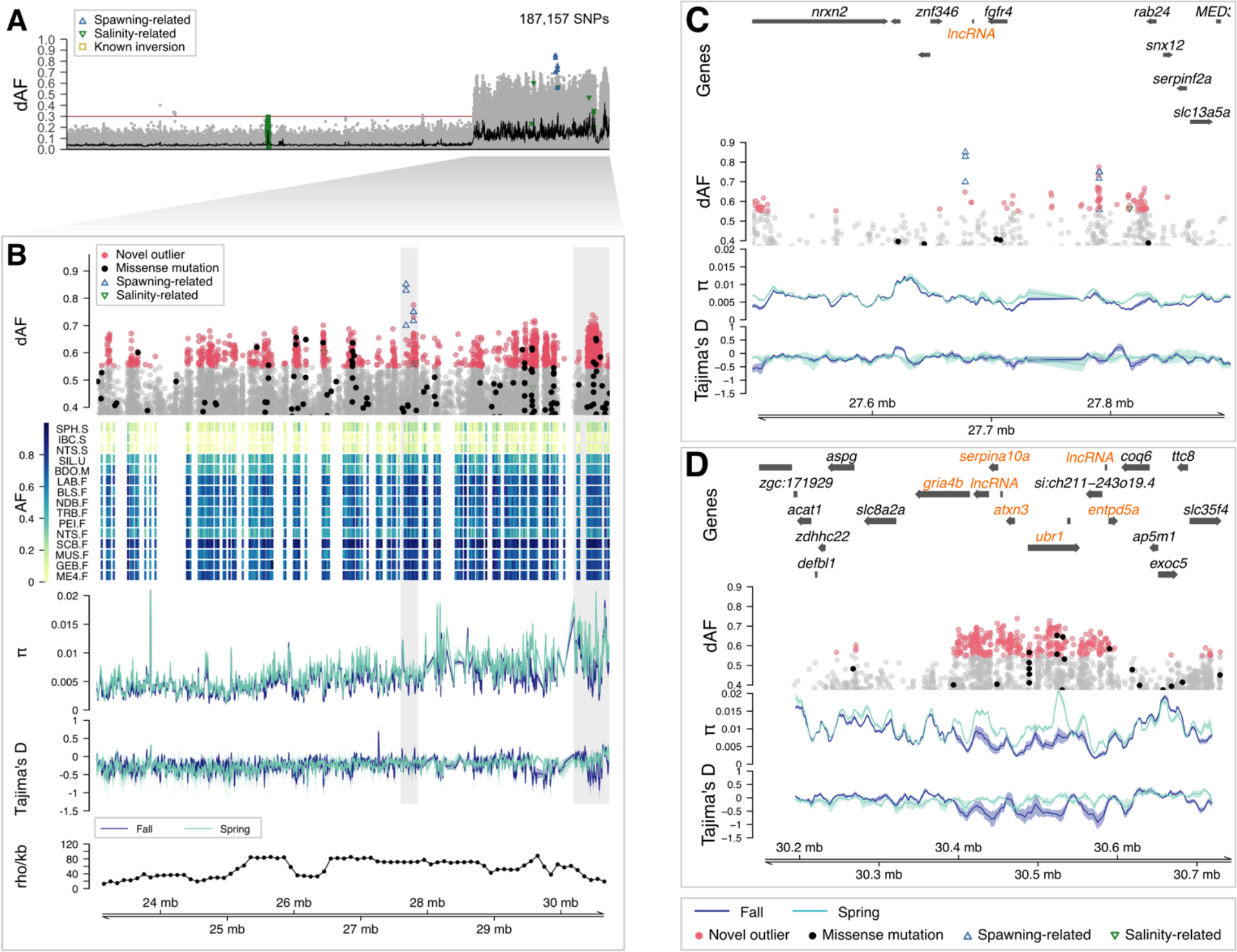
Novel putative inversion on chromosome 8 associated with seasonal reproduction. (**A**) Genetic differentiation (dAF) between spring and fall spawners across chromosome 8. Each dot is a single SNP. The black line is the rolling average of dAF over 100 SNPs, and the horizontal red line is the Bonferroni significance value. SNPs with known association with spawning, salinity, or four known inversions (in chr 6, 12, 17, and 23) (Han et al., 2020) are denoted with empty blue upward triangles, green downward triangles, and yellow squares, respectively. (**B**) Close-up plot to the putative inversion. This plot has five tracks, which show from top to bottom: genetic differentiation between spring and fall spawners for SNPs with dAF ≥ 0.4; heatmap plot depicting the minor allele frequency per population (rows) for the novel outlier SNPs (columns); average nucleotide diversity (π) and Tajima’s D (window size 10 Kbp, step size 2 Kbp) for spring and fall spawners, in light and dark blue lines, respectively; and estimate of recombination rate (rho/Kbp) every 100 Kbp (Pettersson et al., 2019). Novel outlier SNPs (dAF ≥ 0.55) are denoted as red filled circles, missense mutations as filled black circles, and other SNPs are gray circles. Zoom-in into two informative intervals, (**C**) chr8:27,500,000-27,900,000, which contains the most differentiated SNPs, and (**D**) chr8:30,150,000-30,750,000. Genes in the associated region are highlighted in orange.

**Figure 4.**
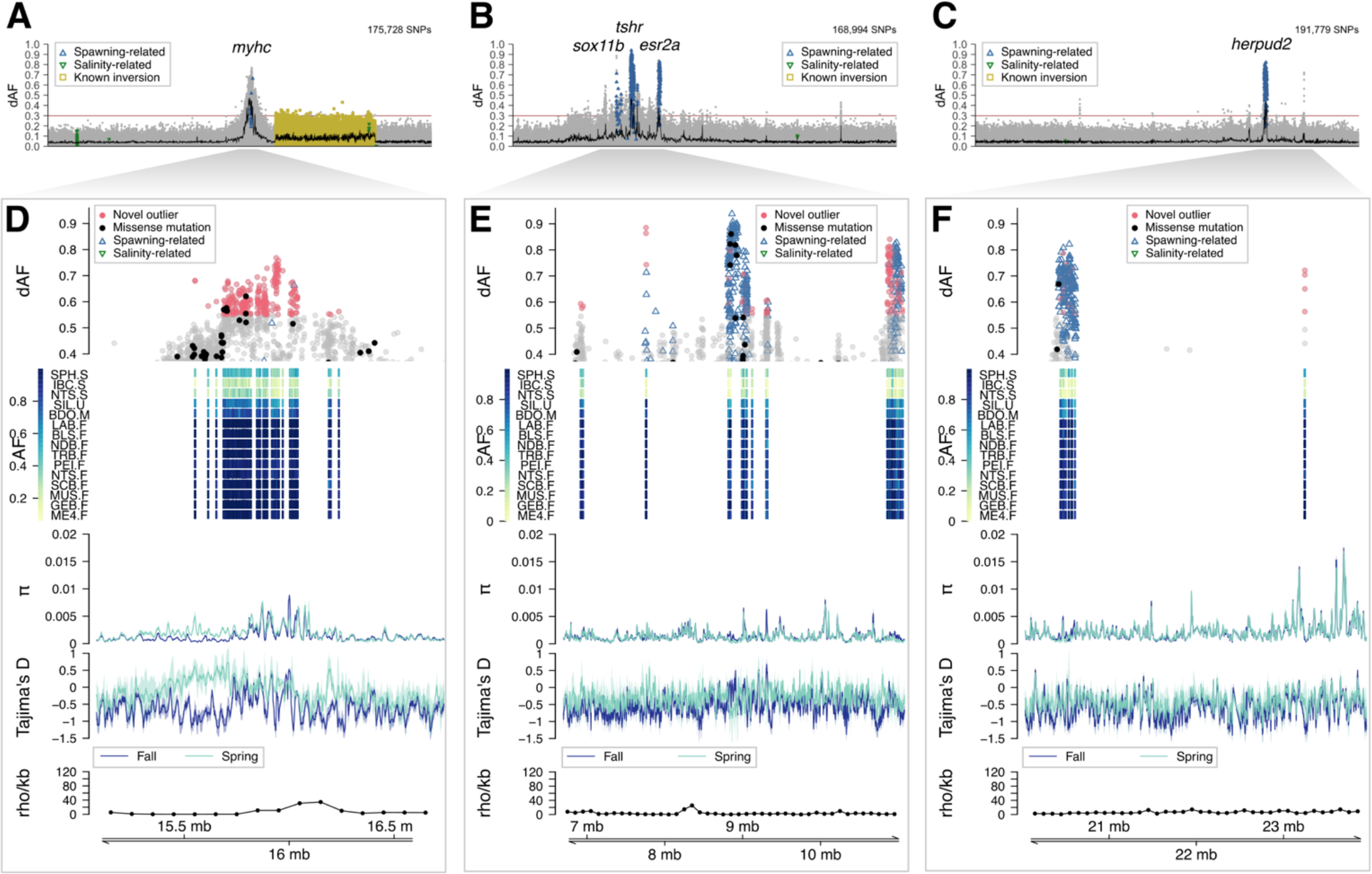
Signatures of selection associated with seasonal reproduction. Genetic differentiation (dAF) between spring and fall spawners across chromosomes (**A**) 12, (**B**) 15, and (**C**) 19. Each dot represents a single SNP. The rolling average of dAF over 100 SNPs is shown as a horizontal black line, and the Bonferroni significance value is shown as a horizontal red line. SNPs with described association with spawning, salinity, or four inversions (in chr 6, 12, 17, and 23) in Han et al. (2020) are indicated with empty blue upward triangles, green downward triangles, and yellow squares, respectively. Close-up to the signatures of selection in chromosomes (**D**) 12, (**E**) 15, (**F**) 19. Each plot has five tracks that show from top to bottom: genetic differentiation between spring and fall spawners for SNPs with dAF ≥ 0.4; heatmap plot depicting the minor allele frequency per population (rows) for the novel outlier SNPs (columns); average nucleotide diversity (π) and Tajima’s D (window size 10 Kbp, step size 2 Kbp) for spring and fall spawners, in light and dark blue lines, respectively; and estimate of recombination rate (rho/Kbp) every 100 Kbp (Pettersson et al., 2019). Novel outlier SNPs (dAF ≥ 0.55) are shown as red filled circles, and missense mutations, as filled black circles. Other SNPs are shown in gray. For zoom-in plots to each peak of divergence including gene names see Supplementary Figures S8, S9, S11.

Our data also revealed that the latitudinal genetic pattern discriminating populations that spawn north or south of a biogeographic transition zone on the eastern Scotian Shelf (Figure 1A) is underlined by a large 8-Mbp long SV. This locus corresponds to a known chromosomal inversion on chr 12 (n = 1106 SNPs with dAF ≥ 0.42, Figure 2B) (Pettersson et al., 2019).

### 3.4 A candidate inversion on chr 8

The new putative SV on chr 8 is located towards one end of the chromosome, in a region with heterogeneous rate of recombination (Figure 3). The genetic differentiation at this locus shows the typical profile of a chromosomal inversion, “block” shaped with sharp “edges” defined by a drastic drop in dAF (Figure 3A) implying reduced recombination. The majority of outlier SNPs at this locus are novel, except for six known spawning time-and two salinity-associated markers (Han et al., 2020) (Figure 3B). This putative inversion was not highlighted in Han et al. (2020), but an examination of the patterns of segregation shows that its variant haplotype is present in some populations in the northeast Atlantic as well (Figure S7).

Alternative haplotypes of the inversion differentiate most spring and fall spawning populations, and intermediate allele frequencies were observed in the summer spawning population from Sept Îsles (SIL-U), the mixed sample from Bras D’Or lake (BDO-M), and the fall spawning population from Northumberland Strait (NTS-F). A comparison of allele frequencies between west and east Atlantic herring revealed that while opposite alleles distinguish spring and fall spawners among Canadian samples, this was not always the case for other Atlantic and Baltic populations (Figure S7A). For instance, the haplotype predominant among Canadian fall spawners is also in high frequency in the summer-spawning sample from Greenland and in two fall-spawning populations from the northwest of England (Orkney, Cape Wrath, Hebrides, Figure S7B). This haplotype is in intermediate frequency in Iceland and in both, spring and fall spawners in the North Sea. The opposite haplotype is prevalent among most spring and winter spawners across the Atlantic, but in the Baltic Sea there is a recombinant haplotype that is shared between autumn spawners and the Pacific herring.

The most differentiated SNPs were located in the interval chr8:27,500,000-27,900,000 (Figure 3C), which harbors some novel and previously reported SNPs. Genes at this locus include a lncRNA (long-non-coding RNA) and *fgfr4* (fibroblast growth factor receptor 4). Another interesting region is located in the interval chr8:30,150,000-30,750,000 (Figure 3D), which shows a significant drop in nucleotide diversity and negative Tajima’s D among fall spawners. This locus harbors several genes, two of which include missense mutations, *ubr1* (ubiquitin protein ligase E3 component n-recognin 1) and *entpd5a* (ectonucleoside triphosphate/diphosphohydrolase 5). A description of outlier SNPs and the closest genes at divergent genomic regions can be found in Table S4.

### 3.5 Novel and previously reported signals of selection

The selection signal on chr 12 is located upstream of a previously described inversion (Pettersson et al., 2019), and extends over 1 Mbp towards the middle of the chromosome in a region of low recombination (Figure 4A, 4D, S8A). Notably, most outlier SNPs in this locus are novel, except for two spawning time-associated markers (Figure 4D, first track, S8B), but they occur in a region showing strong signal of differentiation in previous studies (Han et al., 2020). Alternative alleles were close to fixation between spring and fall spawners, except in the spring-spawning sample from Stephenville (SPH-S, southwest of Newfoundland) that shows intermediate allele frequencies (Figure 4B, second track, S8B). The profile of genetic diversity metrics for the most divergent SNPs revealed that this region likely experienced a selective sweep in the fall spawners, supported by the lower nucleotide diversity and more negative Tajima’s D than in the spring spawners (Figure 4D, S8B). Genes in this region include: five genes of the myosin heavy chain family (i.e., *myhc*-like, *myhz1.1* and three *myh2*), *scd5* (stearoyl-CoA desaturase 5), *tmem150c* (transmembrane protein 150C), *sbno1* (strawberry Notch Homolog 1), and *denr* (density-regulated protein). Opposite alleles at this locus distinguish Canadian and Baltic spring and fall spawners, but this is not applicable to all north Atlantic populations (Figure S8). The “fall” Canadian allele is prevalent in all other oceanic populations (Greenland, Ireland-UK, North Sea, and Norwegian fjords) regardless of their spawning season, and in the Baltic fall spawners. There is a gradient of allele frequencies in the transition zone between the Baltic Sea and the NE Atlantic Ocean.

The signal of selection on chr 15 is 4 Mbp long (Figure 4B, 4E, S9) and it is located near the middle of the chromosome in a region of low recombination. It consists of five distinct loci, two more than detected in the east Atlantic (Han et al., 2020). In genomic order, the first locus is novel, it is approximately 250 Kbp long (chr15:6,750,000-7,000,000), and it harbours the genes *rab15* (member RAS Oncogene Family 15) and *sipa1l1* (signal induced proliferation associated 1 like 1) (Figure S9C). The second locus extends over 200 Kbp (chr15:7,650,000-7,850,000) and contains the gene *sox11b* (Figure S9D), which has been associated with spawning time before (Han et al., 2020) but the two most differentiated SNPs are novel (dAF > 0.7), as they did not show consistent association between northeast Atlantic and Baltic populations. The third locus is about 530 Kbp long (chr15:8,540,000-9,070,000) and harbours a large number of SNPs and genes with a known association with spawning time such as *tshr* and *calm1b* (Figure S9E). The nucleotide diversity and Tajima’s D profiles support a selective sweep in the *tshr* region, in concordance with previous studies (Chen et al., 2021) (Figure S9E). The fourth locus is novel, extends over 150 Kbp (chr15:9,200,000-9,350,000), and contains two genes, *flrt2* (fibronectin leucine-rich transmembrane protein 2) and a *lncRNA* (Figure S9F). The fifth locus extends 280 Kbp (chr15:10,820,000-11,100,000) and consists of two peaks, one spans two well-characterized genes, *esr2a* and *syne2* (Figure S9G), and the other one encompasses several *lncRNA* genes and *l3hypdh* (trans-L-3-hydroxyproline dehydratase). Interestingly, most SNPs in the *esr2a*-*syne2* region are novel, while the SNPs in the *lncRNAs*-*l3hypdh* region have been previously associated with spawning time (Han et al., 2020). Across all five loci in this region, the spring and fall spawners from Canada have opposite alleles in high frequency (Figure S9B, second track). When comparing allele frequencies between west and east Atlantic herring, a similar pattern distinguishing spring and fall spawners is observed in three of the five loci, harbouring *tshr*, *sox11b*, and *flrt2-lncRNA* genes, and in the second peak of the *esr2a*-*syne2-lncRNAs*-*l3hypdh* locus (*lncRNAs*-*l3hypdh*) (Figure S10C, S10E). In the first peak of the latter locus, corresponding to *esr2a*-*syne2*, the “fall” Canadian allele is predominant among populations from the transition zone and the Baltic Sea (Figure S10C, S10D). In the *rab15-sipa1l1* locus, the alternative alleles appear to be unique of Canadian spring spawners (Figure S10A, S10B).

The selection signal on chromosome 19 is located towards one end of the chromosome, in a region of low recombination, and it contains two loci (Figure 4C, 4F, S11). The first locus is 410 Kbp long (19:20,290,000-20,700,000) and harbors the genes *herpud2*, *adgrb1b* and a *lncRNA* (Figure S11C). Most SNPs and genes at this locus have been previously associated with spawning time (Han et al., 2020). The second locus is novel, extends over 145 Kbp (19:23,155,000-23,300,000), and includes a *lncRNA* and *sgk-like* genes. All the SNPs in this locus are novel (Figure S11B). The low nucleotide diversity and negative Tajima’s D profiles at this region suggest that it constitutes a selective sweep (Figure S11D). Indeed, the variant alleles at this locus are unique for the Canadian fall spawners and the summer-spawning sample from Greenland (Figure 11E, S11F).

Overall, this study revealed 10 genomic regions showing highly significant genetic differentiation among the population samples from the northwest Atlantic. Only two of these loci are unique to the northwest Atlantic, and do not appear to be segregating in northeast populations of Atlantic herring (Figure 5, Table S6). These are the loci harboring *rab15-sipa1l1* on chromosome 15 (Figure S9C) and the most distal region on chromosome 19 harboring *lncRNA-sgk-like* (Figure S11D). At some of the shared loci between northwest and northeast populations, like *tshr* and *herpud2* (Figure S9E and S11C, respectively), the alternate haplotypes segregating at these loci are nearly identical and the ranking of top SNPs are almost identical in northwest and northeast herring (Figure 5B, Table S6). In contrast, at some other shared loci there is a clear difference in haplotype composition (Figure 5C). The best example is the region containing a cluster of myosin heavy chain genes on chromosome 12 where numerous SNPs show strong genetic differentiation in northwest Atlantic but low differentiation in northeast populations, whereas a smaller set of other SNPs show the opposite trend suggesting that different haplotypes must be favored by selection in the two sides of the Atlantic (Figure 5C, S8).

**Figure 5.**
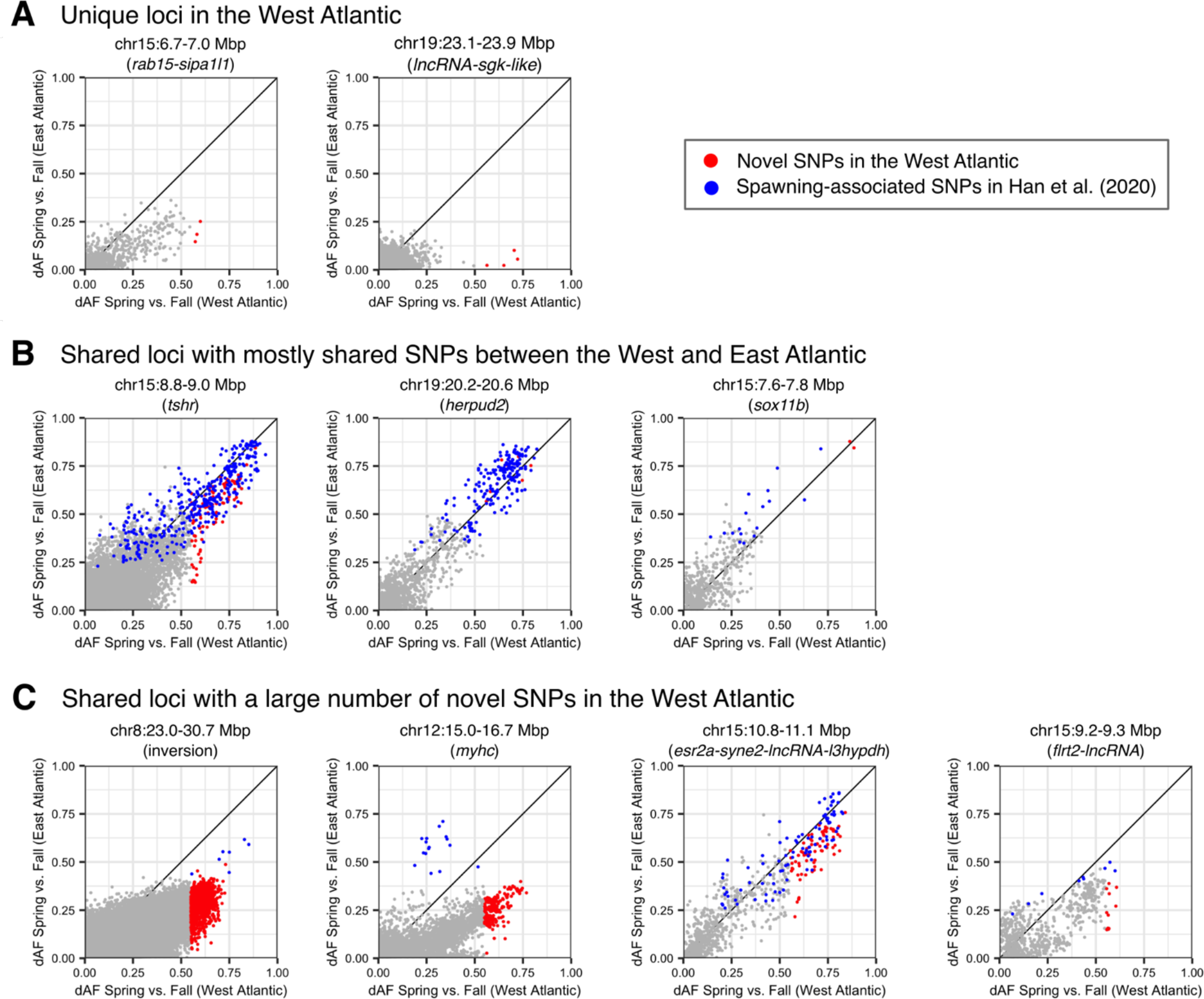
Comparison of the absolute difference in allele frequencies (dAF) between Spring and Fall spawners in the West and East Atlantic for nine genomic regions strongly associated with spawning time. (**A**) Unique loci in the West Atlantic. (**B**) Shared loci with mostly shared SNPs between the West and East Atlantic. (**C**) Shared loci with a large number of novel SNPs in the West Atlantic. Each dot represents a single SNP. Red dots indicate novel SNPs in the West Atlantic and blue dots indicate SNPs with known association with spawning time as reported in Han et al. (2020). The rest of SNPs in the region (not outliers, dAF in the West Atlantic <= 0.55) are shown in gray. A summary of the percentage of unique and shared loci and SNPs is shown in Table S6.

### 3.6 The inversion on chromosome 12 likely associated with minimal temperature survival

The putative inversion on chr 12 that underlies the north-south genetic pattern contains a large number of outlier SNPs as previously reported (Han et al., 2020), but some of the most differentiated ones are novel for the west Atlantic populations (Figure 6A, 6B). The “north” haplotype of the inversion, as defined in Han et al. (2020), is prevalent among northern samples from Labrador (LAB-F), Newfoundland (NDB-F, TRB-F, SPH-S), Gulf of St. Lawrence (BLS-F, SIL-U, PEI-F, NTS-S), Bras D’Or lake (BDO-M), and Scots Bay in the Bay of Fundy (SCB-F) (Figure S12). The “south” haplotype is in high frequency in the sample from the Gulf of Maine (ME4-F), the southernmost location included in this study. Intermediate haplotype frequencies were common in two samples from the Scotian Shelf (MUS-F, GEB-F) (Figure 6B, S12A).

**Figure 6.**
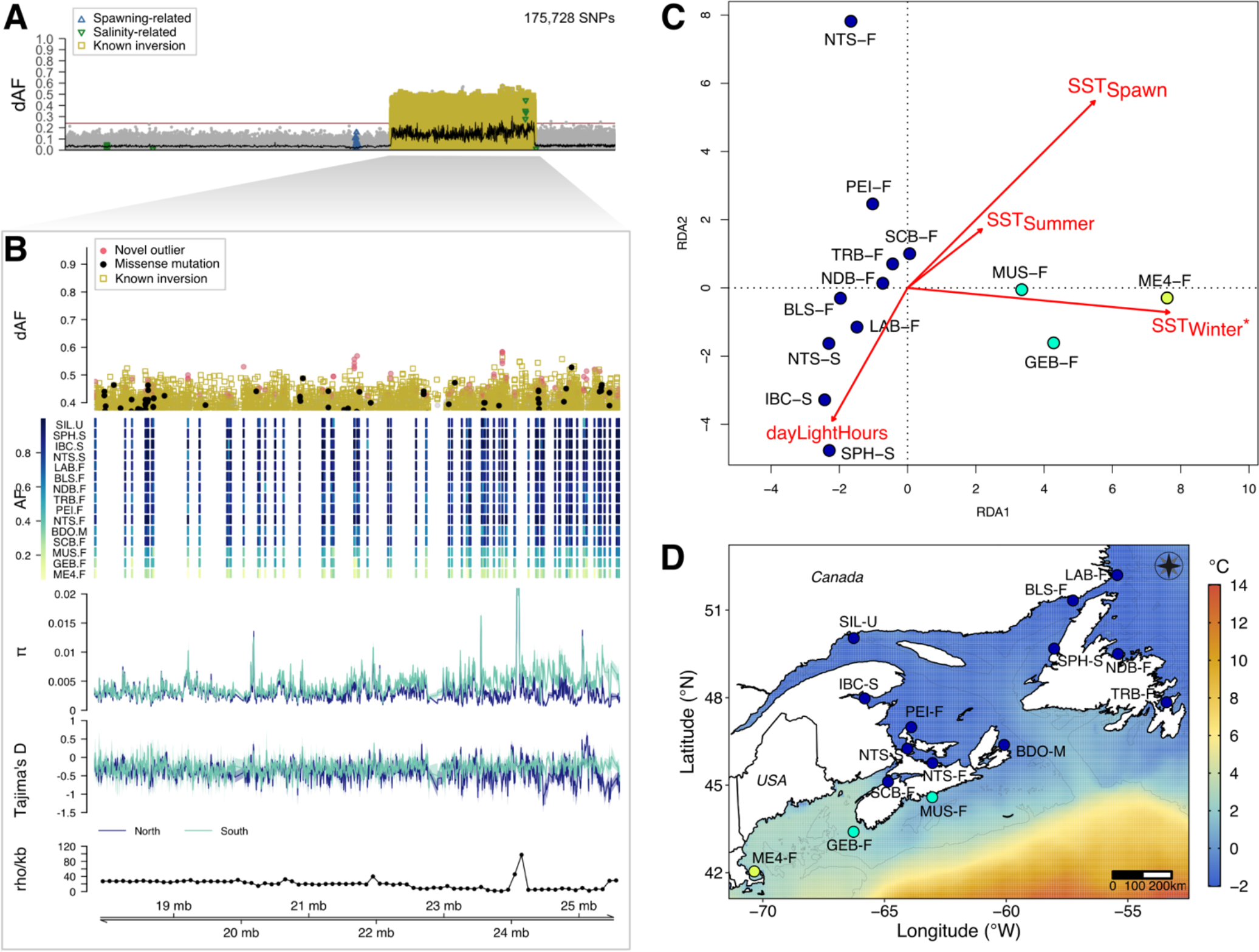
Chromosomal inversion on chromosome 12 associated with spatial genetic divergence along a latitudinal environmental cline. (**A**) Genetic differentiation (dAF) along the chromosome. (**B**) Close-up plot to the putative inversion (chr12:17826318-25603093) consisting of five tracks. The first track is the genetic differentiation for SNPs with dAF ≥ 0.4. Each dot is a single SNP, its shape indicates whether it is a novel outlier (red filled circle), a missense mutation (black circle), or is in one of four inversions reported in Han et al. (2020). The second track depicts the pool-minor allele frequency of the novel outlier SNPs, where each row is a single pool sample, and each column is a SNP. The third and fourth tracks show the profile of the average nucleotide diversity (π) and Tajima’s D of spring and fall spawners, calculated in 10 Kbp sliding windows with a 2 Kbp step. The fifth track shows the recombination rate (rho/Kbp) every 100 Kbp (Pettersson et al., 2019). (**C**) Redundancy analysis plot representing the association between four uncorrelated environmental variables and population allele frequencies of the top outlier SNPs within the inversion on chr 12 (dAF > 0.55). Each circle corresponds to a spawning aggregation and their color indicates their predominant population allele frequency. Sample abbreviations and names as in Table 1. The red arrows (and their length) indicate the level of correlation of each environmental variable with genetic variation in the first two axes. Results of ANOVA to test statistical significance are shown in Table S5. (**D**) Map depicting average winter sea surface temperature and the predominant population allele frequencies at diagnostic SNPs within the inversion on chr 12 for the 15 spawning aggregations included in this study. Each circle corresponds to a spawning aggregation and their color indicates their predominant population allele frequency as per in B) second track.

The current hypothesis is that this inversion is likely associated with adaptation to temperature during spawning (Pettersson et al., 2019). To test this hypothesis, we performed a Genome-Environment Association (GEA) analysis based on redundancy analysis (RDA). We included as environmental variables the number of hours of day light, average sea surface temperature and salinity during the spawning months (SST_Spawn_) and for the winter (SST_Winter_) and summer (SST_Summer_) seasons, as a proxy for the most extreme annual climatic conditions. This analysis indicated that sea surface winter temperature (SST_Winter_) is the environmental factor that best explains the genetic variation of outlier loci in the chr 12 inversion (*P*-value = 0.005, alpha = 0.01) (Figure 6C, S13). Indeed, this was the only variable that was statistically significant for an alpha level of 0.01 (Table S5), and salinity measures were not informative (Figure S14). In the RDA plot, spawning aggregations were separated according to SST_Winter_ along RDA 1, axis that explained 58.1% of the total genetic variance. Notably, the “north” haplotype of the inversion is prevalent among populations that seem to be exposed to much colder temperatures during the winter than the southern ones in the Scotian Shelf and the Gulf of Maine (Figure 6D).

Latitudinal genetic clines can result from natural selection or historical demographic processes, such secondary contact of allopatric populations that diverged during a geologic event and aligned with a contemporary gradient. When natural selection is acting on populations, it would be expected that the fraction of the genome under selection shows the clinal pattern. In contrast, in a scenario where historical divergence leads to limited gene flow, the entire genome should show the cline and it would resemble an isolation-by-distance (IBD) pattern. To examine whether the genetic pattern of the chr 12 inversion followed IBD, we performed a Mantel test in two separate datasets, one consisting of all SNPs and another one of only outlier SNPs in the chr 12 inversion. The test based on all SNPs disclosed that there is no linear relationship between geographic and genetic distances genome-wide (*R*^2^ = 0.04, *P* = 0.11, 9999 replicates). The test performed solely on the SNPs showing the latitudinal genetic cline suggested weak IBD (*R*^2^ = 0.30, *P* = 6e-04, 9999 replicates) and sub-structuring (Figure S14). Taken together, these results suggest that demographic processes do not seem to explain the pattern of latitudinal divergence, although it could coincide with IBD to some extent.

## 4. DISCUSSION

### Population structure only at loci under selection

We found that population structure of NW Atlantic herring mostly lies in a number of genomic regions of adaptive importance, in otherwise weakly differentiated genomes (mean pairwise-*F*_ST_ between 0.001 and 0.018) (Figures 1 and 2). This observation is in agreement with previous studies on Atlantic herring (Bekkevold et al., 2015; Gaggiotti et al., 2009; Guo et al., 2015; Han et al., 2020; Lamichhaney et al., 2012; Limborg et al., 2012; Martinez Barrio et al., 2016; Teacher et al., 2013). The population structure observed is mainly driven by natural selection acting in contrasting environments varying between seasons (spring and fall) and geographic regions (northern and southern) where reproduction and early life stages occur. Our study indicates that the strength of selection must be high, such that adaptive genetic variants display striking allele frequency differences between large populations (in the order of hundreds of thousands to millions of individuals). While herring population structure associated with seasonal reproduction has been reported previously (Han et al., 2020; Lamichhaney et al., 2012; Lamichhaney et al., 2017; Martinez Barrio et al., 2016), our study reveals spatial substructure in the northwest Atlantic that had not yet been identified. Taken together, this work contributes to the growing body of evidence supporting the hypothesis that population structure in marine species is often adaptive and underlies ecological diversity (Benestan et al., 2015; Bradbury et al., 2013; Han et al., 2020; Kess et al., 2021; Lehnert et al., 2018).

This study demonstrates that the majority of loci (8 out of 10) under strong selection in northwest Atlantic herring are also major loci underlying genetic differentiation among populations of herring in the northeast Atlantic. However, there is no perfect overlap regarding which SNPs at these loci show the strongest genetic differentiation between populations. This could be explained by (i) genetic drift due to isolation by distance or by (ii) local genetic adaptation. We conclude that the local adaptation is the most likely explanation because drift should affect all loci in a similar way but this is not the case. We find that the pattern of genetic differentiation is remarkably similar at some loci (in particular *tshr* and *herpud2* both strongly associated with spawning time) whereas other loci, in particular the cluster of myosin heavy chain genes on chromosome 12, show considerable heterogeneity between the west and the east (Figure 5). This result is consistent with our previous conclusion that genetic adaptation in Atlantic herring is to a large extent based on ancestral haplotype blocks, some associated with inversions, that have persisted over hundreds of thousand years (Han et al., 2020; Lamichhaney et al., 2017; Pettersson et al., 2019). However, these haplotype blocks are not static over time, they continue to accumulate causal changes contributing to genetic adaptation, as observed in this study. This is a mode of evolution also noted in the Darwin’s finches and in other adaptive radiations (Rubin et al., 2022).

### Genomic regions associated with adaptation to temporal environmental variation

Populations that predominantly spawn in spring or fall seasons differ at nine major loci distributed in four chromosomes: 8 (one locus), 12 (one locus), 15 (five loci), and 19 (two loci) (Figure 2A, 3, 4). These loci vary in size, ranging from a few Kbp up to several Mbp, as it is the case of the 8-Mbp long structural variant (SV) on chr 8, first described here for the species (Figure 3). This SV is also in high frequency in the sample from Greenland and among fall-spawning oceanic populations. A recombinant variant of the putative inversion is present in Baltic fall-spawning samples (Figure S7). This SV contains numerous genes, for which it is hard to infer its function without further experiments. A region within the SV (chr8:30,150,000- 30,750,000) showing low nucleotide diversity and negative Tajima’s D (Figure 3B, 3D), harbors genes that contain missense mutations, *ubr1* (ubiquitin protein ligase E3 component n-recognin 1) and *entpd5a* (ectonucleoside triphosphate/diphosphohydrolase 5). *ubr1* modulates digestive organ development (He et al., 2017), and *entpd5a* plays a critical role in bone formation as well as in the regulation of phosphate homeostasis (Huitema et al., 2012), both in zebrafish.

Signatures of selection on chr 12 (one locus), chr 15 (five loci), and chr 19 (two loci) encompass both known and novel genetic variants and candidate genes (Figure 4, S8, S9, S11) with respect to those previously associated with ecological adaptation (Han et al., 2020). In agreement with previous research (Han et al., 2020; Lamichhaney et al., 2017; Martinez Barrio et al., 2016), we found that several genetic variants in genes such as *tshr, herpud2, sox11b, esr2a* and *syne2* are shared between NW and NE Atlantic populations. This result confirms that these genes are strong candidates to be involved in seasonal reproduction in Atlantic herring, consistent with the fact that several of them have a known function in reproduction in birds and mammals (Bondesson et al., 2015; Melamed et al., 2012; Ono et al., 2008). Moreover, we discovered novel genetic variants, in particular, in *myhc* in-tandem copies (i.e., *myhc*-like, *myhz1.1* and *myh2*), which are involved in myogenesis (Figure 4D), and in *esr2a* (estrogen receptor beta), which is essential for female reproduction (Figure 4E) (Lu et al., 2017). Interestingly, muscle development and estrogen action are strongly affected by temperature and/or photoperiod (Jin et al., 2010; Johnston et al., 2001), two of the most contrasting environmental conditions between spring and fall seasons. Overall, the spring season is characterized by colder temperatures and increasing daylength in contrast to the fall season, which is characterized by relatively warmer temperatures and decreasing daylength. Considering the potential role of these genes and the presence of genetic variants with particular allele frequency patterns and some seemingly private (Figure S11), we infer that herring populations are locally adapted to the environmental conditions in the NW Atlantic.

Local adaptation to the NW Atlantic is further supported by the discovery of new loci and candidate genes associated with ecological adaptation, with potential important roles in embryological development and gene expression regulation (Figure S8, S9, S11). For example, on chr 12, *sbno1* (strawberry notch homolog 1) is a gene required for normal development of the zebrafish central nervous system (Takano et al., 2010), and *denr* (density-regulated protein) is a highly conserved gene involved in translation initiation (Skabkin et al., 2010). In chr15, *rab15* (member RAS oncogene family 15) presumably participates in cellular response to insulin stimulus and protein metabolism (Bradford et al., 2022), *sipa1l1* (signal induced proliferation associated 1 like 1) is likely involved in vertebrate embryogenesis (Tsai et al., 2007), and *flrt2* (fibronectin leucine-rich transmembrane protein 2) regulates embryonic heart morphogenesis in mice (Müller et al., 2011). In chr19, a long-non coding RNA (lncRNA) gene likely participates in the regulation of gene expression and is characteristic of the NW Atlantic. Indeed, lncRNA genes were often present in several of the genomic regions under selection (Figure 3C, 3D, S8G, S8F, S11C, S11D). lncRNA genes encode RNA molecules longer than 200 bp that are not translated into protein. They are involved in gene expression regulation by directly intervening in chromatin remodeling, transcriptional activation, transcriptional interference, RNA processing, and mRNA translation (Statello et al., 2021; Zhang et al., 2019). Thus, an interesting venue for future research could focus on understanding the role of lncRNAs in local adaptation of Atlantic herring.

### Genomic regions associated with adaptation to spatial environmental variation

Fall spawning populations breeding north or south of the biogeographic transition zone on the Scotian shelf (∼44.61°N) (Stanley et al., 2018), differ in genetic variants spanning a large single locus on chr 12 that corresponds to a known 7.8-Mbp long chromosomal inversion (Figure 6). Spring spawning populations were restricted to the Gulf of St Lawrence, an area north of the transition zone. All spring and fall spawning populations north of the transition zone had the “north” haplotype in high frequency, the “south” haplotype was prevalent in the southernmost location in Maine (ME4-F), and populations south of the transition zone in Musquodoboit harbor (MUS-F) and German Banks (GEB-F) had intermediate frequencies (Figure 6A, 6B).

While the role of this inversion is unknown, it has been proposed that it is associated with adaptation to temperature at spawning (Pettersson et al., 2019). Our Genome-Environment Association analysis indicated that minimal temperature during winter is the most concordant environmental factor with the spatial pattern of genomic differentiation driven by the putative inversion on chromosome 12 (Figure 6C, S10, S11) (*P* = 0.005, alpha = 0.01). The strong thermal gradient in this area results in those populations north of the biogeographic break experiencing significantly colder temperatures on average than those south of the break (Figure 6D). This pattern agrees with other studies comparing genetic and environmental gradients in this area (Lehnert et al., 2018; Stanley et al., 2018). Temperature is a critical factor influencing fish physiology (Reynolds & Casterlin, 1980) and distribution (García Molinos et al., 2016). Temperature has a particularly strong influence on the physiology of early life stages, which, due physiological constrains and high mortality during this phase, are particularly sensitive to thermal conditions (Dahlke et al., 2020; Marr, 1956). Correspondingly, our results suggest that selection associated with early life survival during the coldest months of the year (i.e., post-settlement mortality) may underlie patterns of genetic structure. Connectivity during the early life history of herring in the NW Atlantic has been generally assumed to be limited, corresponding to fine spatial scales of population structure (Stephenson et al., 2009). Our results correspond to and further illustrate this hypothesis, whereby emergent genetic patterns associated with adult fish correspond to processes likely imparting selection on earlier life history phases.

The prevalence of intermediate frequencies near the transition zone suggests that climatic conditions may vary between years in relation to oceanographic regional trends (Townsend et al., 2004). In a study of genetic structure of the invasive European green crab (*Carcinus meanus*), Jeffery et al. (2018) similarly identified intermediate populations proximate to the biogeographic break within our focal area. It is possible that populations at these locations experience significant inter-annual environmental fluctuations during winter, depending on the strength of the warm Gulf Stream flowing north or of the cold Labrador Current flowing south. Thus, we infer that balancing selection may be favoring both inversion haplotypes at locations near the transition zone.

### Implications

The results of this work have numerous implications in evolutionary biology and fisheries management of Atlantic herring, and for other marine species with comparable life histories. This study expands our understanding of the genomic basis of adaptation with gene flow by: (i) contributing a list of candidate genetic variants and genes underlying intraspecific diversity; (ii) providing genetic evidence supporting the hypothesis that strong selection can maintain molecular divergence at loci underlying variation of phenotypic traits and fitness in large populations despite the presence of gene flow; (iii) illustrating how such selection can be driven by ecological and environmental factors varying at different temporal and spatial scales; and (iv) confirming the central role of structural variants in adaptation with gene flow. This knowledge is important to improve our ability to predict adaptive responses of wild populations in a scenario of global warming. This study also illustrates the complexity predicting how climate change will influence marine populations that exhibit fine-scale, environmentally mediated population structure, whereby changes in temperature could elicit asymmetric responses among population subgroups and potentially disrupting synchrony between reproductive timing, larval emergence and seasonal prey availability (sensu (Cushing, 1990) – ‘Match-Mismatch hypothesis’).

Our results highlight the genetic underpinnings of the fine-scale population structure that has long-been associated with herring in the NW Atlantic (Stephenson et al., 2009). In a period of declining abundance, a robust assessment of genetic differentiation provides critical information for management to conserve intraspecific variation. As we now have the molecular basis to distinguish various herring ecotypes, a subset of outlier SNP markers described here can be used for assessing stock composition. Such approach could help to monitor mixed stock fishing amongst management regions helping to minimize the risk of overexploitation of vulnerable components (Bekkevold et al., 2015; Hemmer-Hansen et al., 2019; Larson et al., 2014).

## Acknowledgements

We thank staff at Fisheries and Oceans Canada, the Maine Department of Resources, Comeau’s Seafoods Ltd, Cape Breeze Seafoods Ltd, and fishers Crystal and Donald Kent, Gordon McKay, and many others for their valuable contribution in sample and tissue collection. Thanks to Gregory McCracken for assistance in sampling and DNA extraction, to Gavin Douglas, Emma Sylvester, Sarah Lehnert, Simone Fior, Alan Bergland, Miguel Carneiro, Michael Blum, and Mathieu Gautier for data analysis assistance. We thank Zeliang Wang and David Brickman from the Bedford Institute of Oceanography for access to BNAM model outputs. Thanks to Sean Rogers and Rob Beiko for constructive comments that helped improve this manuscript. Thanks to Luis Fuentes, Edith Pardo, Gloria Fuentes, Scott Campbell, Cathy Campbell, Meagan Campbell, John Sherlock, and the Ruzzante Lab for enriching discussions and support. Computations were mainly carried out on the supercomputer Mp2 of the University of Sherbrooke, managed by Calcul Québec and Compute Canada and funded by the Canada Foundation for Innovation (CFI), the ministère de l’Économie, de la science et de l’innovation du Québec (MESI) and the Fonds de recherche du Québec - Nature et technologies (FRQ-NT). The analyses comparing northeast and northwest Atlantic populations were performed using the computational infrastructure provided by the Swedish National Infrastructure for Computing (SNIC) at UPPMAX partially funded by the Swedish Research Council through grant agreement no. 2018-05973. A.P.F.P. and D.E.R. thank the Killam Trust. A.P.F.P. thanks the Vanier Canada Graduate Scholarship, the President’s Award of Dalhousie University, the Nova Scotia Graduate Scholarship, the Lett Fund and a Strategic grant to D.E.R for graduate studies funding. This study was funded by NSERC Discovery and Strategic grants to D.E.R.

## Dat1a Archiving Statement

Sequence data generated in this study is available in the BioProject PRJNA930418 of the NCBI. Oceanographic data per location used in GEA and population-allele frequencies per SNP and pool are available in Dryad *(code will be provided upon manuscript approval)*. Custom scripts are available in the Github repository: https://github.com/apfuentes/2023_NWAtlanticHerring_PopGen.

