## Supplementary Information for "Adaptation to seasonal reproduction and temperature-associated factors drive temporal and spatial differentiation in northwest Atlantic herring despite gene flow"

**Table of Contents:**


| | |
|---|---|
| **Table S1** | Page 2 |
| **Table S2** | Page 2 |
| **Table S3** | Page 3 |
| **Table S4** | Page 4 |
| **Table S5** | Page 4-5 |
| **Table S6** | Page 6 |
| **Figure S1** | Page 7 |
| **Figure S2** | Page 8 |
| **Figure S3** | Page 9 |
| **Figure S4** | Page 10 |
| **Figure S5** | Page 11 |
| **Figure S6** | Page 12 |
| **Figure S7** | Page 13 |
| **Figure S8** | Page 14 |
| **Figure S9** | Page 16 |
| **Figure S10** | Page 18 |
| **Figure S11** | Page 19 |
| **Figure S12** | Page 21 |
| **Figure S13** | Page 22 |
| **Figure S14** | Page 22 |
| **Figure S15** | Page 23 |
| **Supplementary references** | Page 24 |

**Table S1.** Read mapping summary statistics of the pool-seq data of 15 herring spawning aggregations included in this study.

| Locality | Code | Total reads (in millions) | GC (%) | Insert size (in bp) | Median coverage | Mean coverage | Aligned reads (%) | Sequence batch |
|---|---|---|---|---|---|---|---|---|
| Sept Îles | SIL-U | 478.9 | 43 | 546 | 66.0X | 20.6X | 96.20 | 2016 |
| Inner Baie Des Chaleurs | IBC-S | 434.5 | 43 | 478 | 57.0X | 17.7X | 96.20 | 2015 |
| Stephenville | SPH-S | 510.3 | 43 | 505 | 72.0X | 22.8X | 96.70 | 2016 |
| Northumberland Strait | NTS-S | 426.6 | 44 | 523 | 58.0X | 19.3X | 96.10 | 2015 |
| Northumberland Strait | NTS-F | 431.9 | 43 | 504 | 57.0X | 17.7X | 95.50 | 2015 |
| Labrador | LAB-F | 474.7 | 43 | 534 | 69.0X | 22.4X | 98.50 | 2016 |
| Blanc Sablon | BLS-F | 487.9 | 43 | 531 | 67.0X | 20.6X | 96.00 | 2016 |
| Notre Dame Bay | NDB-F | 472.4 | 43 | 534 | 69.0X | 22.5X | 98.90 | 2016 |
| Trinity Bay | TRB-F | 453.5 | 43 | 503 | 63.0X | 20.0X | 96.60 | 2015 |
| Prince Edward Island | PEI-F | 479.7 | 43 | 536 | 69.0X | 22.5X | 98.10 | 2016 |
| Bras D'Or lake | BDO-M | 488.9 | 43 | 526 | 68.0X | 21.7X | 96.30 | 2016 |
| Scots Bays | SCB-F | 502.5 | 43 | 539 | 73.0X | 23.5X | 98.80 | 2016 |
| Musquodoboit | MUS-F | 470.7 | 43 | 533 | 68.0X | 21.6X | 98.80 | 2016 |
| German Banks | GEB-F | 424.8 | 43 | 532 | 57.0X | 17.8X | 95.80 | 2015 |
| Maine fishing area 514 | ME4-F | 558.8 | 44 | 469 | 77.0X | 26.9X | 96.80 | 2016 |

**Table S2.** Pairwise $\hat{F}_{ST}^{pool}$ for 15 herring spawning aggregations in the northwest Atlantic.

| | BDO-M | BLS-F | GEB-F | IBC-S | LAB-F | ME4-F | PEI-F | MUS-F | NDB-F | NTS-F | NTS-S | SCB-F | SIL-U | SPH-S | TRB-F |
|---|---|---|---|---|---|---|---|---|---|---|---|---|---|---|---|
| **BDO-M** | 0 | 0.0022 | 0.0024 | 0.006 | 0.0011 | 0.0065 | 0.0015 | 0.001 | 9.00E-04 | 0.0026 | 0.004 | 0.0022 | 0.0062 | 0.0077 | 0.0015 |
| **BLS-F** | 0.0022 | 0 | 0.0037 | 0.0098 | 0.0012 | 0.0086 | 0.002 | 0.0022 | 0.0013 | 0.0035 | 0.0077 | 0.0021 | 0.0077 | 0.0105 | 0.0021 |
| **GEB-F** | 0.0024 | 0.0037 | 0 | 0.0114 | 0.0024 | 0.0049 | 0.003 | 0.001 | 0.0022 | 0.0047 | 0.0094 | 0.0024 | 0.009 | 0.0124 | 0.0026 |
| **IBC-S** | 0.006 | 0.0098 | 0.0114 | 0 | 0.0088 | 0.0164 | 0.0087 | 0.0098 | 0.0083 | 0.0086 | 0.0032 | 0.0113 | 0.0108 | 0.0085 | 0.0087 |
| **LAB-F** | 0.0011 | 0.0012 | 0.0024 | 0.0088 | 0 | 0.0073 | 0.001 | 0.001 | 3.00E-04 | 0.0026 | 0.0068 | 0.001 | 0.0069 | 0.0097 | 0.001 |
| **ME4-F** | 0.0065 | 0.0086 | 0.0049 | 0.0164 | 0.0073 | 0 | 0.0077 | 0.0043 | 0.0068 | 0.0094 | 0.0145 | 0.0068 | 0.0139 | 0.0175 | 0.0072 |
| **PEI-F** | 0.0015 | 0.002 | 0.003 | 0.0087 | 0.001 | 0.0077 | 0 | 0.0016 | 0.001 | 0.0029 | 0.0067 | 0.0018 | 0.0072 | 0.0096 | 0.0016 |
| **MUS-F** | 0.001 | 0.0022 | 0.001 | 0.0098 | 0.001 | 0.0043 | 0.0016 | 0 | 8.00E-04 | 0.0032 | 0.0077 | 0.0013 | 0.0075 | 0.0106 | 0.0014 |
| **NDB-F** | 9.00E-04 | 0.0013 | 0.0022 | 0.0083 | 3.00E-04 | 0.0068 | 0.001 | 8.00E-04 | 0 | 0.0023 | 0.0063 | 0.0012 | 0.0067 | 0.0092 | 0.001 |
| **NTS-F** | 0.0026 | 0.0035 | 0.0047 | 0.0086 | 0.0026 | 0.0094 | 0.0029 | 0.0032 | 0.0023 | 0 | 0.0064 | 0.004 | 0.0081 | 0.0098 | 0.0029 |
| **NTS-S** | 0.004 | 0.0077 | 0.0094 | 0.0032 | 0.0068 | 0.0145 | 0.0067 | 0.0077 | 0.0063 | 0.0064 | 0 | 0.0092 | 0.0089 | 0.0066 | 0.0068 |
| **SCB-F** | 0.0022 | 0.0021 | 0.0024 | 0.0113 | 0.001 | 0.0068 | 0.0018 | 0.0013 | 0.0012 | 0.004 | 0.0092 | 0 | 0.0083 | 0.0118 | 0.0019 |
| **SIL-U** | 0.0062 | 0.0077 | 0.009 | 0.0108 | 0.0069 | 0.0139 | 0.0072 | 0.0075 | 0.0067 | 0.0081 | 0.0089 | 0.0083 | 0 | 0.0125 | 0.0074 |
| **SPH-S** | 0.0077 | 0.0105 | 0.0124 | 0.0085 | 0.0097 | 0.0175 | 0.0096 | 0.0106 | 0.0092 | 0.0098 | 0.0066 | 0.0118 | 0.0125 | 0 | 0.0098 |
| **TRB-F** | 0.0015 | 0.0021 | 0.0026 | 0.0087 | 0.001 | 0.0072 | 0.0016 | 0.0014 | 0.001 | 0.0029 | 0.0068 | 0.0019 | 0.0074 | 0.0098 | 0 |

**Table S3.** Pool data samples used from Han et al (2020), of which 47 correspond to Atlantic herring and one to Pacific herring.

| Sample name | Latitude | Longitude | Region | Spawning season | Sampling date | Sample size | Salinity (ppt) | Reference |
|---|---|---|---|---|---|---|---|---|
| A_Kalix_Baltic_Spring | 65.52 | 22.43 | Baltic Sea | Spring | 19800629 | 47 | 3 | Lamichhaney et al., 2012 |
| HGS1_Riga_Baltic_Spring | 58.34 | 24.62 | Baltic Sea | Spring | 20140421 | 96 | 5.5 | Hill et al. 2019; Bekkevold et al. 2016 |
| HGS2_Riga_Baltic_Spring | 58.34 | 24.62 | Baltic Sea | Spring | 20160530 | 100 | 5.5 | Hill et al. 2019; Bekkevold et al. 2016 |
| HGS3_Riga_Baltic_Autumn | 58.10 | 23.92 | Baltic Sea | Autumn | 20140903 | 96 | 5.5 | Hill et al. 2019; Bekkevold et al. 2016 |
| HGS4_Riga_Baltic_Autumn | 58.10 | 23.92 | Baltic Sea | Autumn | 20150909 | 99 | 5.5 | Hill et al. 2019; Bekkevold et al. 2016 |
| PB6_Gävle_Baltic_Summer | 60.43 | 17.18 | Baltic Sea | Summer | 20120718 | 100 | 6 | Martinez Barrio et al. 2016 |
| B_Vaxholm_Baltic_Spring | 59.26 | 18.18 | Baltic Sea | Spring | 19790827 | 50 | 6 | Lamichhaney et al. 2012 |
| PB1_Hästskär_Baltic_Spring | 60.35 | 17.48 | Baltic Sea | Spring | 20130522 | 50 | 6 | Martinez Barrio et al. 2016 |
| PB4_Hudiksvall_Baltic_Spring | 61.45 | 17.30 | Baltic Sea | Spring | 20120419 | 100 | 6 | Martinez Barrio et al. 2016 |
| PB5_Gävle_Baltic_Spring | 60.43 | 17.18 | Baltic Sea | Spring | 20120507 | 100 | 6 | Martinez Barrio et al. 2016 |
| PB7_Gävle_Baltic_Autumn | 60.44 | 17.35 | Baltic Sea | Autumn | 20120904 | 100 | 6 | Martinez Barrio et al. 2016 |
| G_Gamleby_Baltic_Spring | 57.50 | 16.27 | Baltic Sea | Spring | 19790820 | 49 | 7 | Lamichhaney et al. 2012 |
| PB11_Kalmar_Baltic_Spring | 57.39 | 17.07 | Baltic Sea | Spring | 20120509 | 100 | 7 | Martinez Barrio et al. 2016 |
| PB12_Karlskrona_Baltic_Spring | 56.10 | 15.33 | Baltic Sea | Spring | 20120530 | 100 | 7 | Martinez Barrio et al. 2016 |
| HGS71_Rugen_Baltic_Spring | 54.14 | 13.47 | Baltic Sea - Transision | Spring | 20090406 | 40 | 8 | Hill et al. 2019; Limborg et al. 2012 |
| HGS72_Rugen_Baltic_Spring | 54.14 | 13.47 | Baltic Sea - Transision | Spring | 20030424 | 40 | 8 | Hill et al. 2019; Limborg et al. 2012 |
| PN3_CentralBaltic_Baltic_Spring | 55.24 | 15.51 | Baltic Sea (=Transition gene pool) | Spring | 20111018 | 100 | 8 | Martinez Barrio et al. 2016 |
| S18_Germany_Baltic | 54.22 | 13.58 | Baltic Sea - Transision | Spring | 20120320 | 110 | 8 | Present study |
| HGS12_BornholmBasin_Baltic_Autumn | 55.30 | 15.22 | Baltic Sea - Transision | Autumn | 20161110 | 45 | 8 | Hill et al. 2019 |
| HGS6_Schlei_Baltic_Spring | 54.60 | 9.76 | Baltic Sea - Transision | Spring | 20110509 | 50 | 9 | Hill et al. 2019; Bekkevold et al. 2015 |
| HGS5_Schlei_Baltic_Autumn | 54.60 | 9.76 | Baltic Sea - Transision | Autumn | 20101025 | 89 | 9 | Hill et al. 2019 |
| H_Fehmarn_Baltic_Autumn | 54.50 | 11.30 | Baltic Sea - Transision | Autumn | 19790923 | 50 | 12 | Lamichhaney et al. 2012 |
| HGS11_RingkobingFjord_Spring | 56.02 | 8.19 | Northeast Atlantic Ocean - Transition | Spring | 20090422 | 40 | 12 | Hill et al. 2019; Limborg et al. 2012 |
| HGS24_Landvik_Atlantic_Spring | 58.32 | 8.50 | Northeast Atlantic Ocean - Transition | Spring | 20150428 | 30 | 15 | Present study |
| HGS24_Landvik_Atlantic_Spring | 58.32 | 8.50 | Northeast Atlantic Ocean - Transition | Spring | 20150506 | 16 | 15 | Present study |
| HGS24_Landvik_Atlantic_Spring | 58.32 | 8.50 | Northeast Atlantic Ocean - Transition | Spring | 20150603 | 4 | 15 | Present study |
| LandvikS17_Norway_Atlantic_Spring | 58.32 | 8.50 | Northeast Atlantic Ocean - Transition | Spring | 20150519 | 38 | 15 | Present study |
| J_Träslövslage_Baltic_Spring | 57.03 | 12.11 | Northeast Atlantic Ocean - Transition | Spring | 19781023 | 50 | 20 | Lamichhaney et al. 2012 |
| PB9_Kattegat_Atlantic_Spring | 57.43 | 11.42 | Northeast Atlantic Ocean - Transition | Spring | 20120312 | 100 | 23 | Martinez Barrio et al. 2016 |
| O_Hamburgsund_Atlantic_Spring | 58.30 | 11.13 | Northeast Atlantic Ocean - Transition | Spring | 19790319 | 49 | 25 | Lamichhaney et al. 2012 |
| HGS8_KattegatNorth_Atlantic_Spring | 57.40 | 11.40 | Northeast Atlantic Ocean - Transition | Spring | 20090424 | 41 | 25 | Hill et al. 2019; Limborg et al. 2012 |
| PB10_Skagerrak_Atlantic_Spring | 58.19 | 11.21 | Northeast Atlantic Ocean - Transition | Spring | 20120320 | 100 | 25 | Martinez Barrio et al. 2016 |
| HGS25_Lindås_Atlantic_Spring | 60.73 | 5.13 | Northeast Atlantic Ocean | Spring | 20100312 | 50 | 28 | Present study |
| HGS26_Lusterfjorden_Atlantic_Spring | 61.48 | 7.58 | Northeast Atlantic Ocean | Spring | 20111108 | 50 | 32 | Present study |
| HGS23_Clyde_Atlantic_Spring | 55.14 | -5.04 | Northeast Atlantic Ocean | Spring | 20030314 | 39 | 33 | Present study; Hatfield et al., 2007 |
| HGS17_IsleOfMan_IrishSea_Autumn | 54.06 | -4.37 | Northeast Atlantic Ocean | Autumn | 20150930 | 50 | 33 | Present study |
| HGS19_TeelinBay_Atlantic_Winter | 54.63 | -8.63 | Northeast Atlantic Ocean | Winter | 20160108 | 47 | 34 | Present study |
| HGS10_Downs_EnglishChannel_Winter | 51.34 | 1.90 | Northeast Atlantic Ocean | Winter | 20161212 | 55 | 35 | Present study |
| HGS9_Greenland_Atlantic_Spring | 60.78 | -47.15 | Northwest Atlantic Ocean | Summer | 20161005 | 38 | 35 | Present study |
| HGS15_NSSH_Atlantic_Spring | 67.46 | 9.47 | Northeast Atlantic Ocean | Spring | 20170220 | 43 | 35 | Hill et al. 2019 |
| HGS20_Skye_Atlantic_Spring | 57.41 | -6.13 | Northeast Atlantic Ocean | Spring | 20040224 | 50 | 35 | Present study; Hatfield et al., 2007 |
| HGS27_Gloppen_Atlantic_Spring | 61.77 | 6.16 | Northeast Atlantic Ocean | Spring | 20100802 | 20 | 35 | Present study |
| HGS27_Gloppen_Atlantic_Spring | 61.77 | 6.16 | Northeast Atlantic Ocean | Spring | 20120401 | 11 | 35 | Present study |
| HGS27_Gloppen_Atlantic_Spring | 61.77 | 6.16 | Northeast Atlantic Ocean | Spring | 20120905 | 15 | 35 | Present study |
| HGS27_Gloppen_Atlantic_Spring | 61.77 | 6.16 | Northeast Atlantic Ocean | Spring | 20121130 | 4 | 35 | Present study |
| Q_Norway_Atlantic_Atlantic_Spring | 64.52 | 10.15 | Northeast Atlantic Ocean | Spring | 19800207 | 49 | 35 | Lamichhaney et al. 2012 |
| PB2_Iceland_Atlantic_Spring | 65.49 | -12.58 | Northeast Atlantic Ocean | Spring | 20110915 | 100 | 35 | Martinez Barrio et al. 2016 |
| HGS21_Hebrides_Atlantic_Mixed | 58.17 | -7.23 | Northeast Atlantic Ocean | Mixed | 20160828 | 50 | 35 | Present study |
| HGS18_CelticSea_Atlantic_AutumnWinter | 51.59 | -6.51 | Northeast Atlantic Ocean | Winter | 20151201 | 50 | 35 | Present study |
| HGS16_Orkney_NorthSea_Autumn | 59.00 | -2.00 | Northeast Atlantic Ocean | Autumn | 20150901 | 49 | 35 | Present study |
| HGS22_CapeWrath_Atlantic_Autumn | 58.61 | -4.37 | Northeast Atlantic Ocean | Autumn | 20150910 | 49 | 35 | Present study |
| N_NorthSea_Atlantic_Autumn | 58.06 | 6.10 | Northeast Atlantic Ocean | Autumn | 19790805 | 49 | 35 | Lamichhaney et al. 2012 |
| Pacific herring | | | Pacific Ocean | Spring | 20121124 | 50 | 35 | Martinez Barrio et al. 2016 |

**Table S4.** Outlier SNPs at genomic regions strongly associated with ecological adaptation in NW Atlantic herring (separate Microsoft Excel file).

**Table S5.** RDA model and ANOVA tests of the significance of the RDA model, axes, and environmental variables included in the model.

**RDA model**

Call: rda(formula = poolData_infoReg_final ~ dayLightHours + SST_Summer + SST_Winter + SST_spawn, data = envData_final_sc_set, scale = FALSE)
Inertia Proportion Rank
Total 822.7839 1.0000

Constrained 554.9930 0.6745 4
Unconstrained 267.7908 0.3255 8
Inertia is variance

Eigenvalues for constrained axes:
RDA1 RDA2 RDA3 RDA4
483.2 27.9 25.3 18.6

Eigenvalues for unconstrained axes:
PC1 PC2 PC3 PC4 PC5 PC6 PC7 PC8
100.21 35.92 28.35 25.09 21.08 20.05 19.25 17.83

**Test of significance of the model**

Permutation test for rda under reduced model
Permutation: free
Number of permutations: 999

Model: rda(formula = poolData_infoReg_final ~ dayLightHours + SST_Summer + SST_Winter + SST_spawn, data = envData_final_sc_set, scale = FALSE)
Df Variance F Pr(>F)
Model 4 554.99 4.145 0.003 **
Residual 8 267.79
---
Signif. codes: 0 '***' 0.001 '**' 0.01 '*' 0.05 '.' 0.1 ' ' 1

**Test of significance of the RDA axes**

Permutation test for rda under reduced model
Forward tests for axes
Permutation: free
Number of permutations: 999

Model: rda(formula = poolData_infoReg_final ~ dayLightHours + SST_Summer + SST_Winter + SST_spawn, data = envData_final_sc_set, scale = FALSE)
Df Variance F Pr(>F)
RDA1 1 483.20 14.4350 0.003 **

```
RDA2      1    27.88  0.8328  0.974
RDA3      1    25.27  0.7549  0.950
RDA4      1    18.65  0.5571  0.910
Residual  8   267.79
---
Signif. codes:  0 ‘***’ 0.001 ‘**’ 0.01 ‘*’ 0.05 ‘.’ 0.1 ‘ ’ 1
```

**Test of significance of the environmental variables**

```
Permutation test for rda under reduced model
Marginal effects of terms
Permutation: free
Number of permutations: 1000

Model: rda(formula = poolData_infoReg_final ~ dayLightHours + SST_Summer + SST_Winter + SST_spawn, data
= envData_final_sc_set, scale = FALSE)
Df Variance      F   Pr(>F)
dayLightHours  1   20.739 0.6195 0.521479
SST_Summer     1   26.536 0.7927 0.428571
SST_Winter     1  251.736 7.5204 0.004995 **
SST_spawn      1   25.522 0.7625 0.469530
Residual       8  267.791
---
Signif. codes:  0 ‘***’ 0.001 ‘**’ 0.01 ‘*’ 0.05 ‘.’ 0.1 ‘ ’ 1
```

**Table S6.** Ten genomic regions showing highly significant genetic differentiation among the northwest Atlantic herring populations. Description of the number and percentage of novel and shared SNPs between West and East Atlantic oceanic populations.

| Genomic region | Characteristics | Population contrast | Novel locus | Total number of SNPs | Total number of outlier SNPs* | Number of novel outlier SNPs* | Number of shared+ outlier SNPs* | % of novel SNPs | % of shared SNPs |
|---|---|---|---|---|---|---|---|---|---|
| chr8:23,040,136-30,729,461 | chr8 inversion | Spring vs Fall | No | 102739 | 1356 | 1349 | 7 | 0.995 | 0.005 |
| chr12:15,081,429-16,739,133 | *myhc* | Spring vs Fall | No | 4764 | 270 | 269 | 1 | 0.996 | 0.004 |
| chr15:6,750,000-7,000,000 | *rab15-sipa1l1* | Spring vs Fall | Yes | 854 | 3 | 3 | 0 | 1.000 | 0.000 |
| chr15:7,650,000-7,850,000 | *sox11b* | Spring vs Fall | No | 666 | 5 | 3 | 2 | 0.600 | 0.400 |
| chr15:8,800,000-9,075,000 | *tshr* | Spring vs Fall | No | 868 | 180 | 16 | 164 | 0.089 | 0.911 |
| chr15:9,200,000-9,350,000 | *flrt2-lncRNA* | Spring vs Fall | No | 715 | 15 | 13 | 2 | 0.867 | 0.133 |
| chr15:10,850,000-11,100,000 | *esr2a-syne2-lncRNA-l3hypdh* | Spring vs Fall | No | 943 | 132 | 73 | 59 | 0.553 | 0.447 |
| chr19:20,235,179-20,630,966 | *herpud2* | Spring vs Fall | No | 1534 | 167 | 16 | 151 | 0.096 | 0.904 |
| chr19:23,170,000-23,954,060 | *lncRNA-sgk-like* | Spring vs Fall | Yes | 5968 | 4 | 4 | 0 | 1.000 | 0.000 |
| chr12:17,823,410-25,605,433 | chr12 inversion | Northern vs Southern Fall | No | 64240 | 434 | 51 | 383 | 0.118 | 0.882 |

(*) For the Spring vs Fall contrast, outlier SNPs correspond to those with dAF >= 0.55, and for the Northern vs Southern Fall contrast, outlier SNPs are those with dAF >= 0.45.
(+) Shared SNPs were defined as outlier SNPs with a known association with adaptation as previously reported in Han et al. (2020).

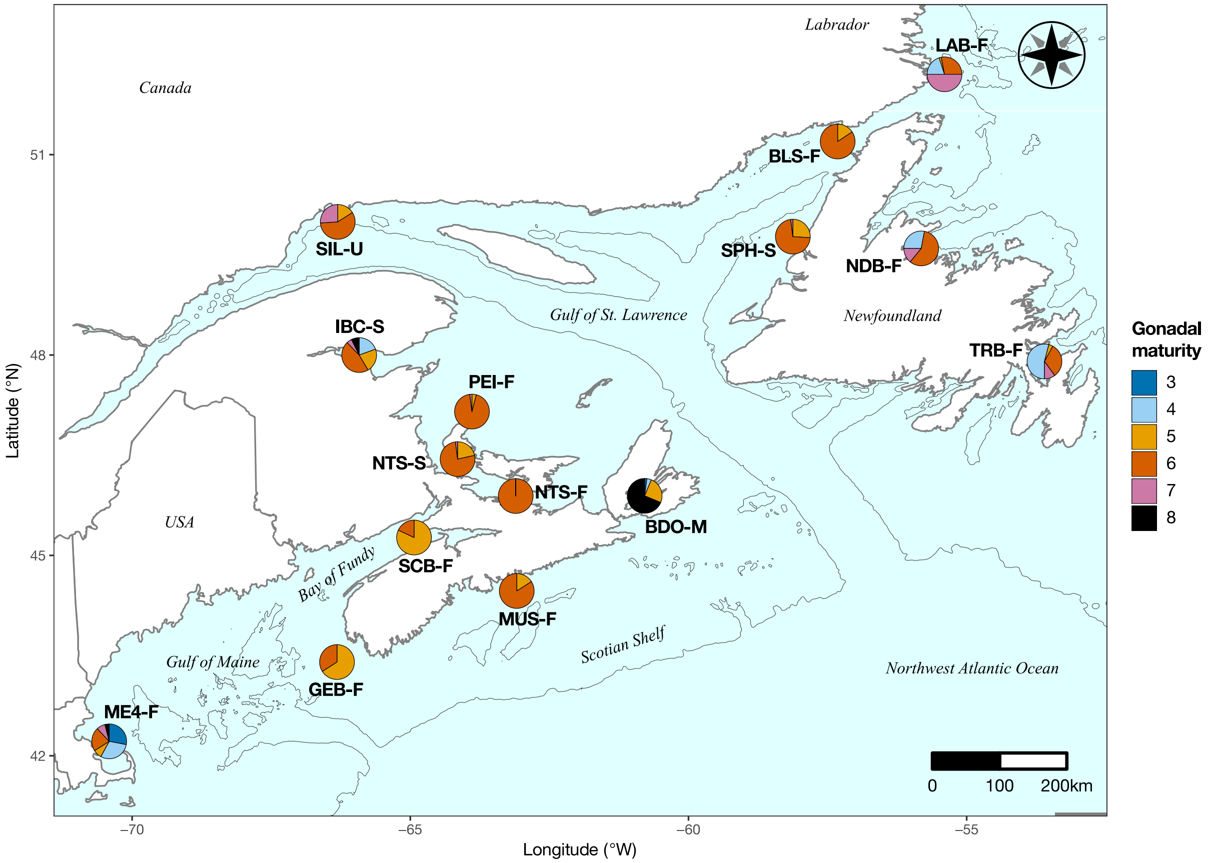


**Figure S1**. Proportion of individuals at various gonadal maturity stages at the time of collection in each of the sampling sites. Gonadal maturity categories: 3 = Mid maturation, 4 = Late maturation, 5 = Spawning capable, 6 = Spawning, 7 = Spent-recovery, 8 = Abnormal (Bucholtz, Tomkiewicz, & Dalskov, 2008).

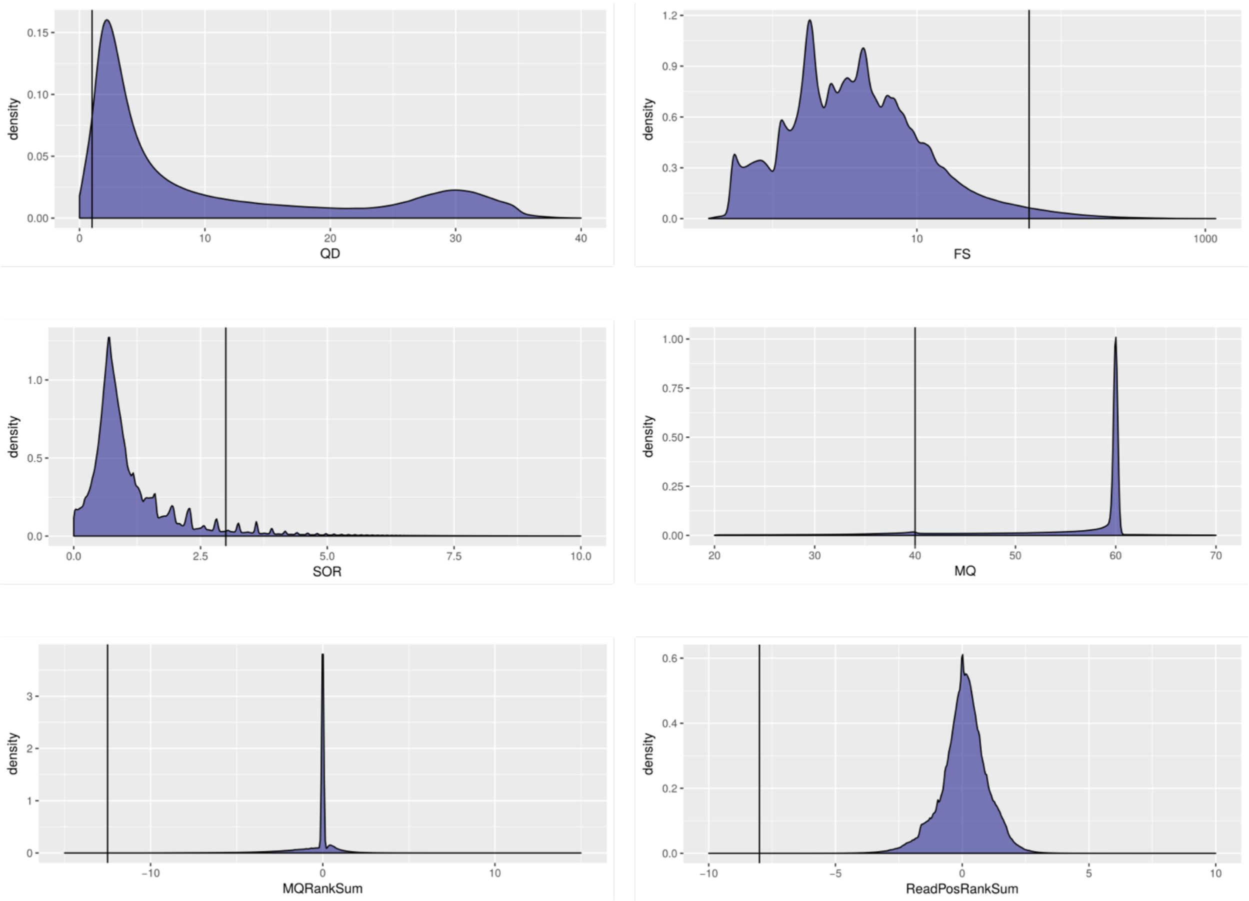


**Figure S2.** Density plots of GATK variant annotations used as reference to determine cutoff values to apply hard filters to raw SNP calls. The black vertical line shows the cutoff value used. Abbreviations: QualByDepth (QD) 2.0, FisherStrand (FS) 60.0, StrandOddsRatio (SOR) 3.0, RMSMappingQuality (MQ) 40.0, MappingQualityRankSumTest (MQRankSum) -12.5, ReadPosRankSumTest (ReadPosRankSum) -8.0. A complete explanation of each of these filters can be found in (Broad Institute, 2016).

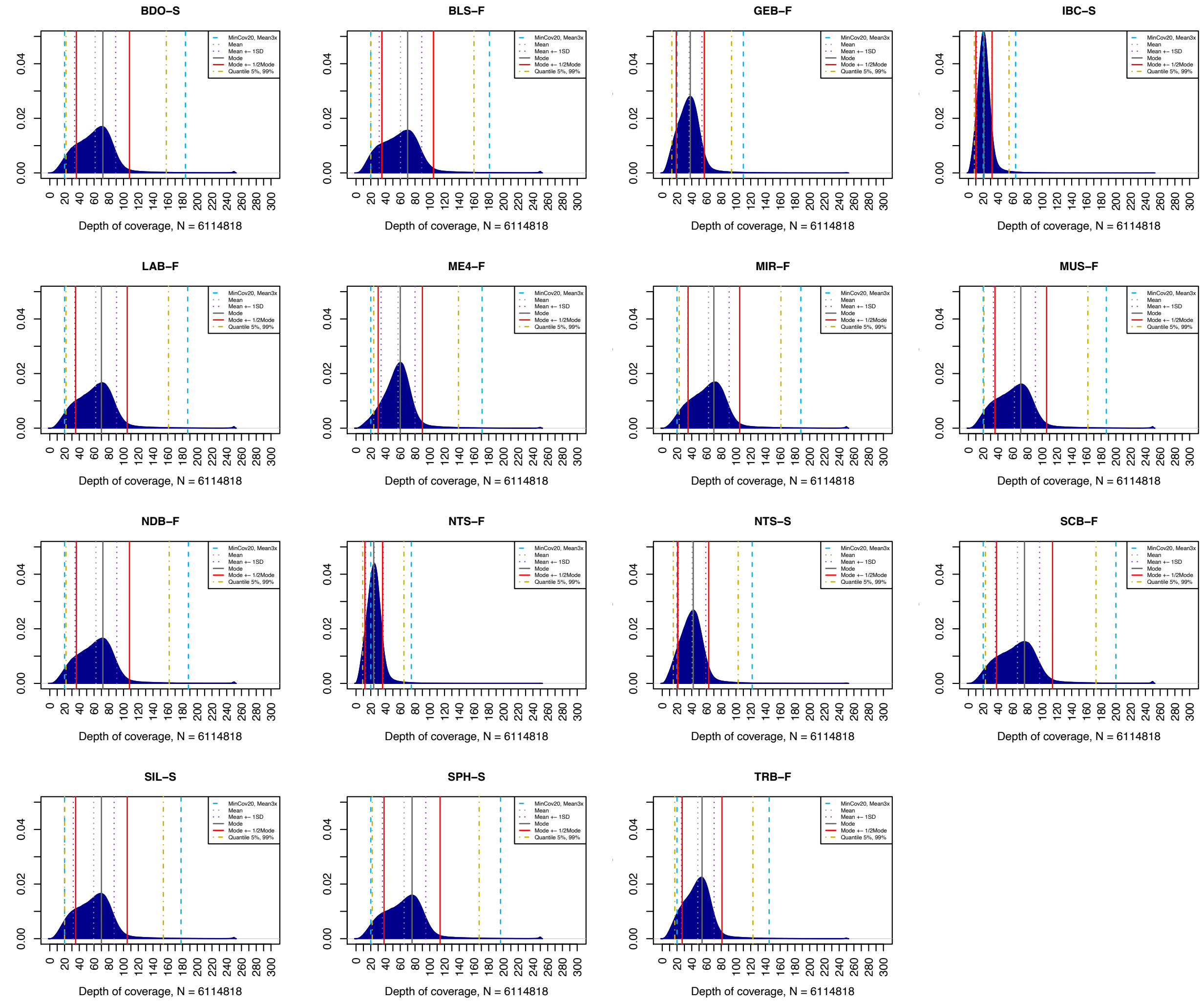


**Figure S3**. Depth of coverage distribution per pool. The vertical lines indicate various measures of central tendency and cut off values. Dashed gray line: mean value, dashed purple line: mean ± 1 standard deviation (SD); gray continuous line: mode, red continuous line: mode ± 1/2 mode; dashed light blue line: minimum coverage 20X and maximum coverage 3 times the mean value; dashed gold line: quantile 5-99%.

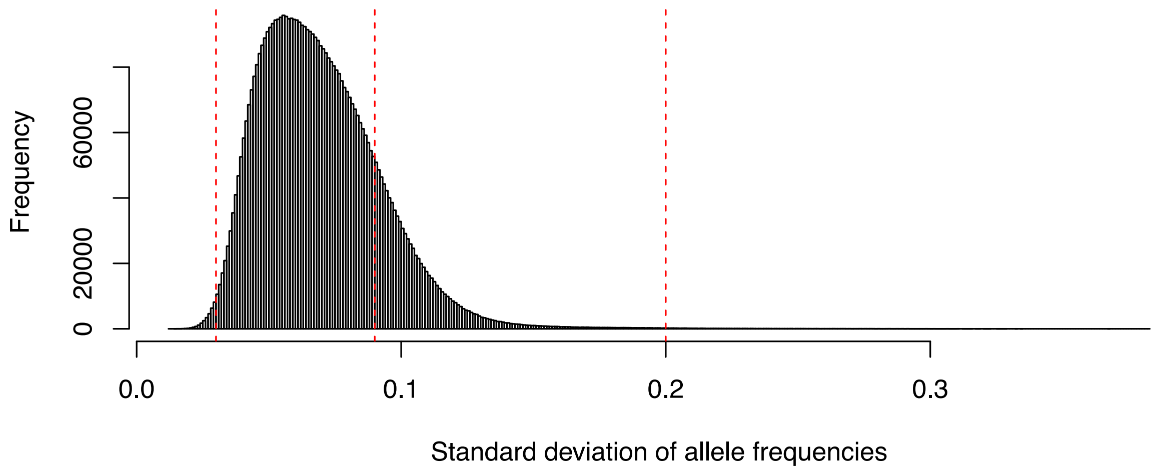


**Figure S4**. Empirical distribution of pool-allele frequencies and they standard deviation. The dashed red lines indicate cut of values to determine the set of undifferentiated markers (with 0.03 < allele frequency SD ≤ 0.09) and highly differentiated markers (allele frequency SD ≥ 0.2 from the mean).

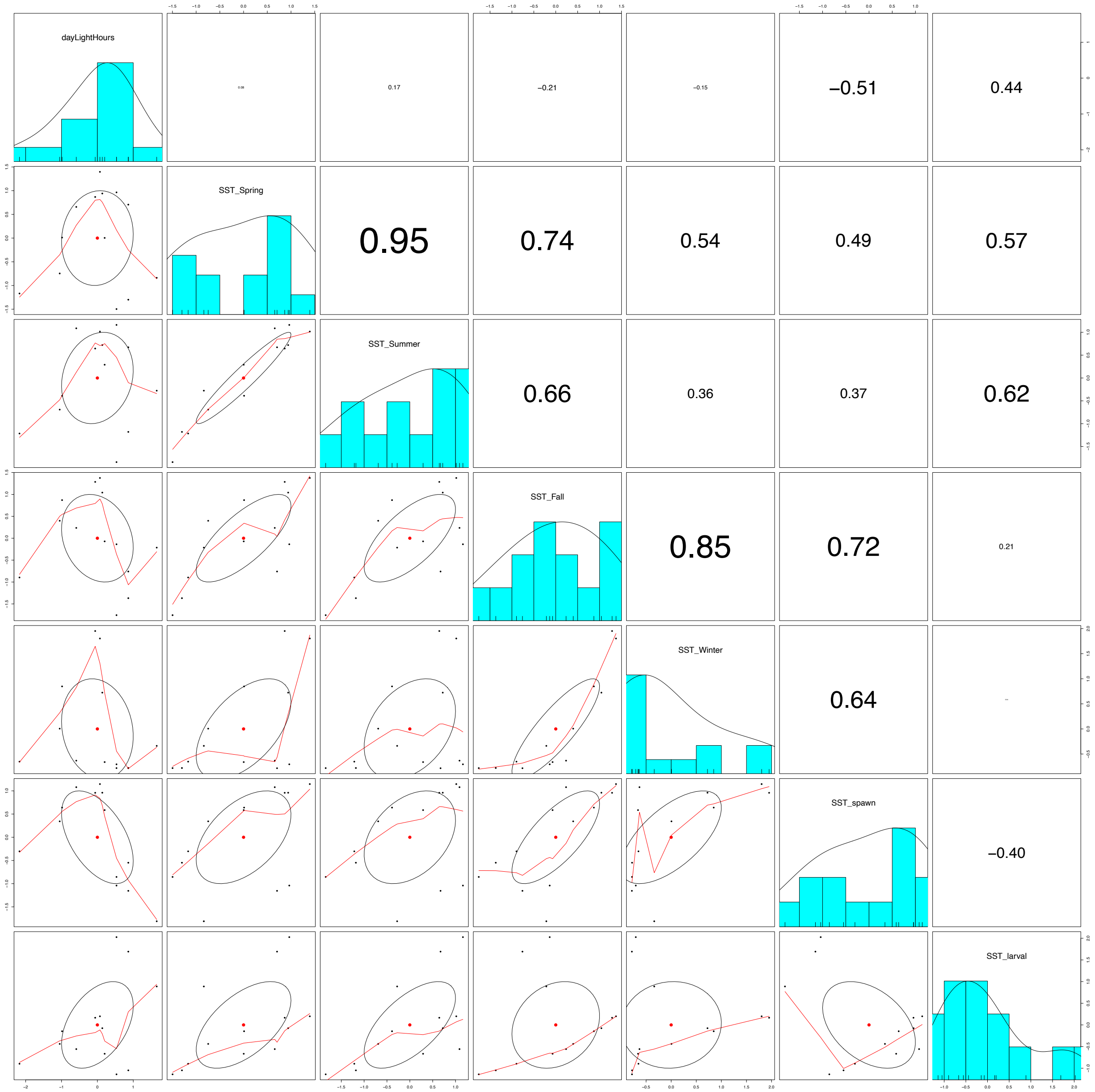


**Figure S5.** Pairwise correlation of the initial set of environmental variables considered for environment association analyses. Below the diagonal a scatterplot for each pairwise comparison is shown, and above the diagonal the correspondent Pearson correlation coefficients are presented; their font size reflects their magnitude.

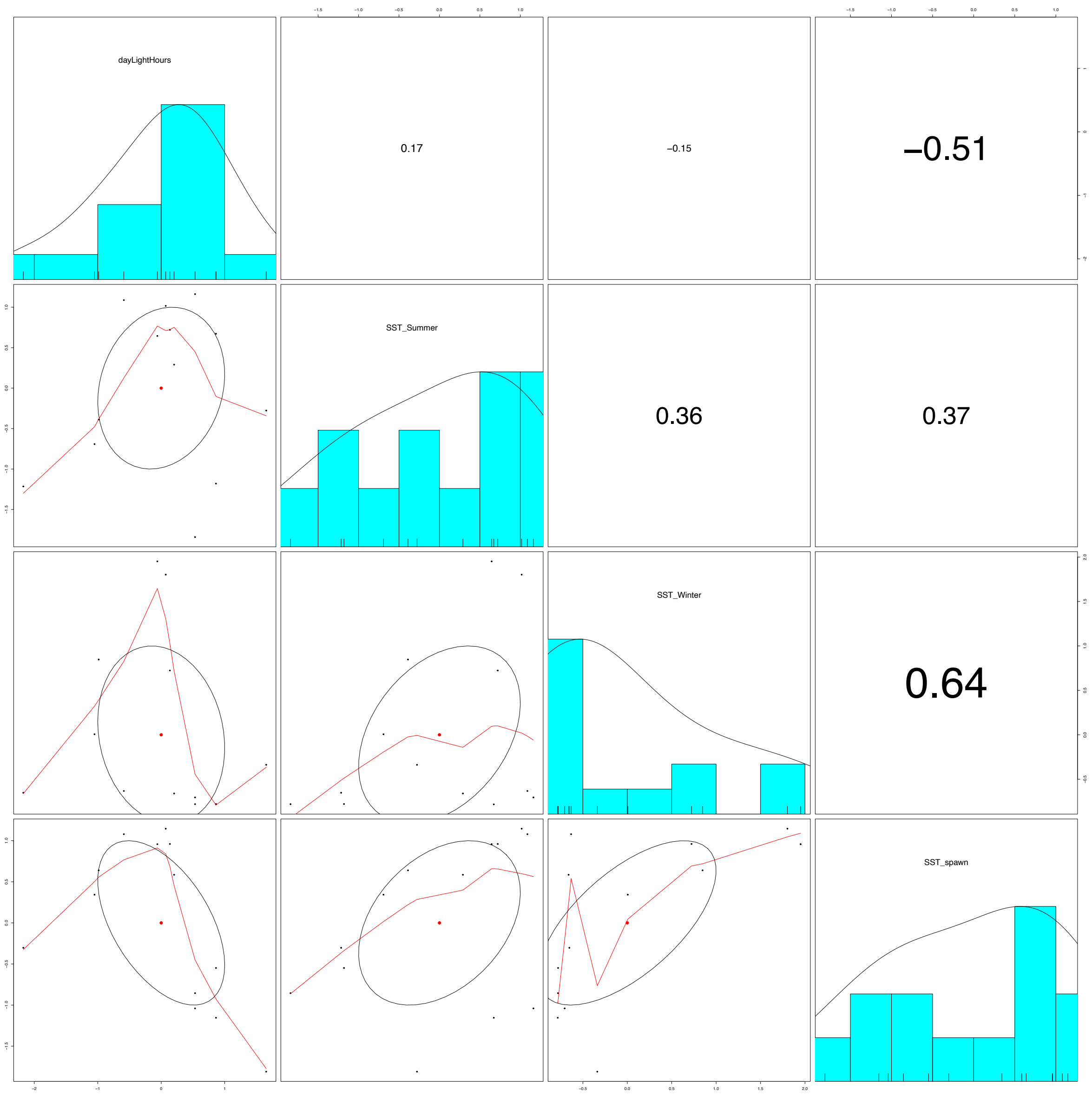


**Figure S6.** Pairwise correlation of the final set of environmental variables considered for environment association analyses. Below the diagonal a scatterplot for each pairwise comparison is shown, and above the diagonal the correspondent Pearson correlation coefficients are presented; their font size reflects their magnitude. The uncorrelated environmental variables were: sea surface summer temperature ($SST_{Summer}$), sea surface winter temperature ($SST_{Winter}$), sea surface temperature at spawning ($SST_{spawn}$), and day light hours (dayLightHours).

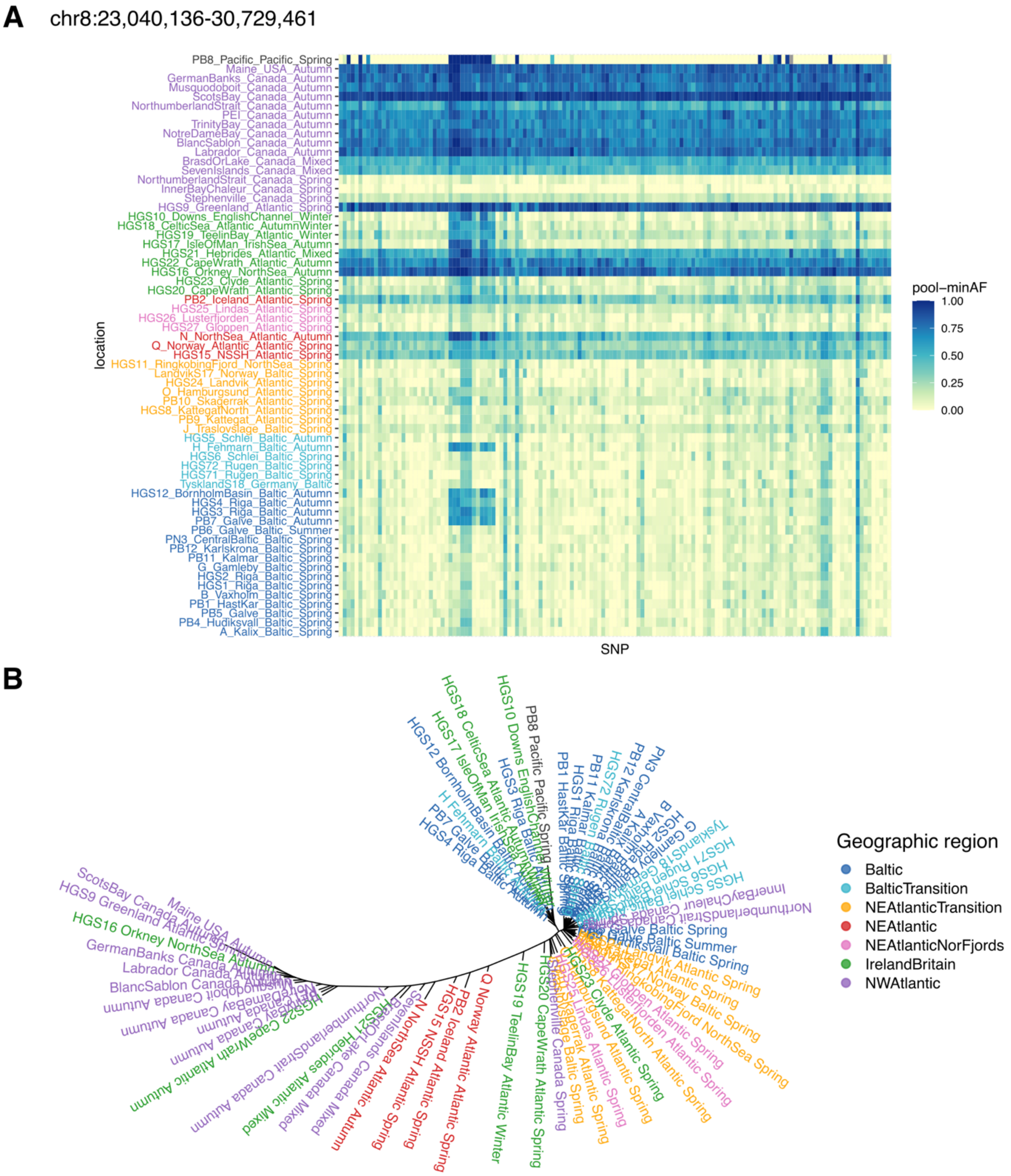


**Figure S7**. Pool-allele frequencies and neighbor-joining tree based on diagnostic variants in the putative inversion on chromosome 8 (dAF >= 0.55).

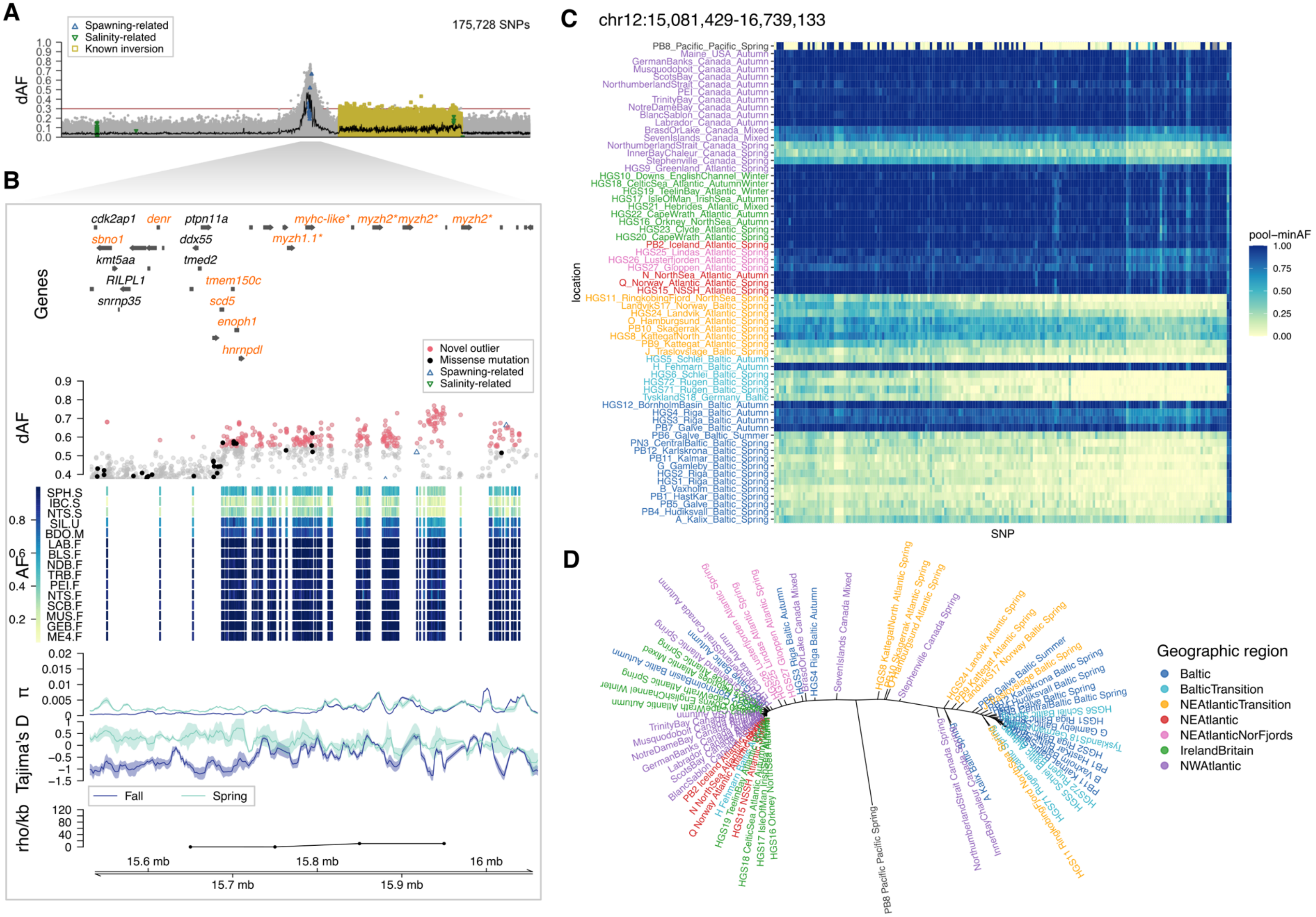


**Figure S8**. Selection signal on chromosome 12. (**A**) Genetic differentiation (dAF) along chr 12. (**B**) Close-up to the target region. This plot consists of five tracks, from top to bottom: gene models; genetic differentiation between spring and fall spawners for SNPs with dAF $\geq$ 0.4. Novel outlier SNPs (dAF $\geq$ 0.55) are denoted as red filled circles, missense mutations as filled black circles, spawning-related SNPs as empty blue triangles, and other SNPs are gray circles; heatmap plot depicting the minor allele frequency per population (rows) for the novel outlier SNPs (columns); average nucleotide diversity ($\pi$) and Tajima's D (window size 10 Kbp, step size 2 Kbp) for spring and fall spawners, in light and dark blue lines, respectively; and estimate of recombination rate (rho/Kbp) every 100 Kbp (Pettersson et al., 2019). (**C**) Pool-allele frequencies and (**D**) neighbor-joining tree based on diagnostic genetic variants (dAF >= 0.55).

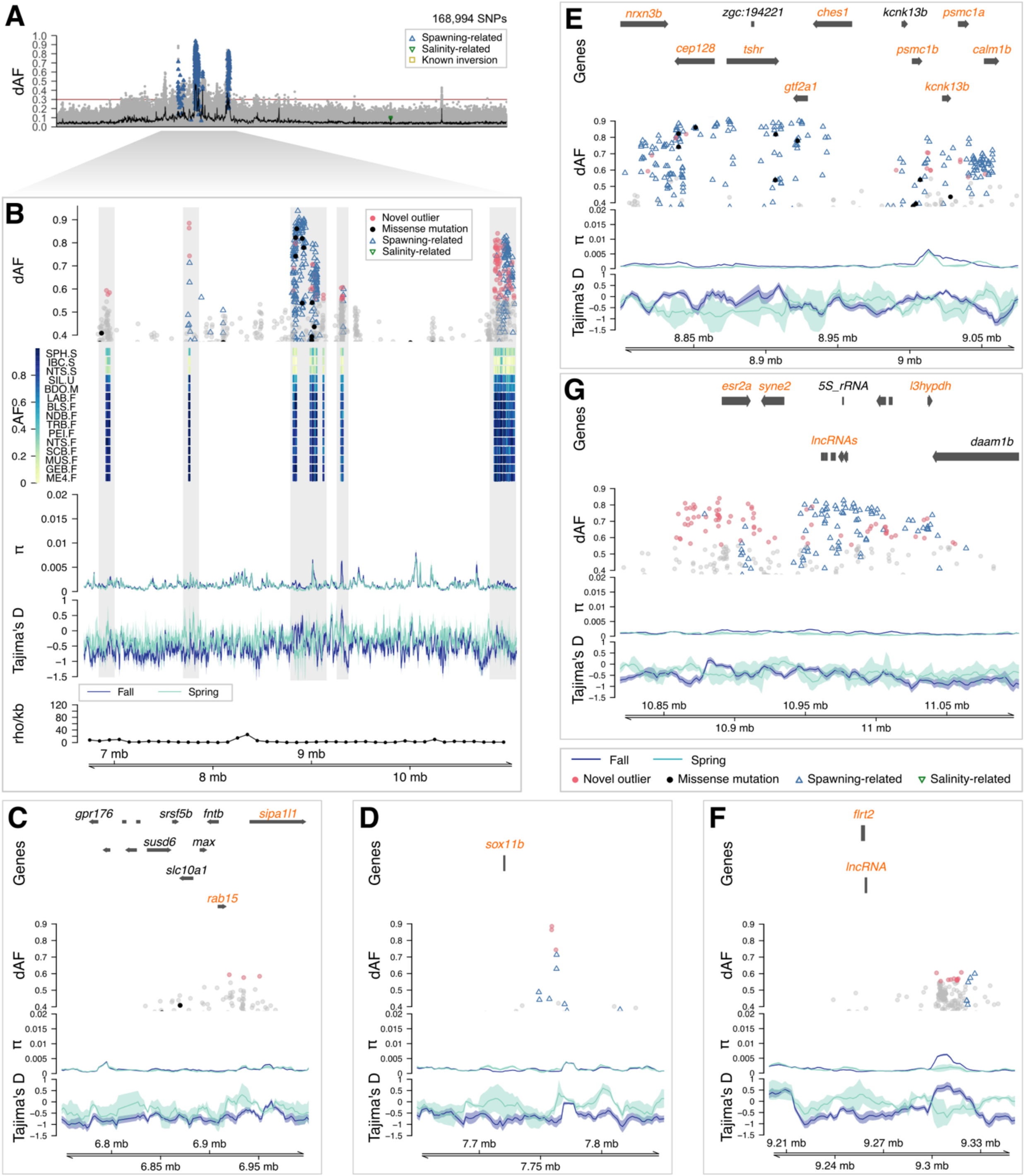


**Figure S9**. Close up to the selection signal on chr 15. (**A**) Genetic differentiation (dAF) along the chromosome. (**B**) Close-up to the target region. This plot consists of five tracks, from top to bottom: genetic differentiation between spring and fall spawners for SNPs with dAF ≥ 0.4. Novel outlier SNPs (dAF ≥ 0.55) are denoted as red filled circles, missense mutations as filled black circles, spawning-related SNPs as empty blue triangles, and other SNPs are gray circles; heatmap plot depicting the minor allele frequency per population (rows) for the novel outlier SNPs (columns); average nucleotide diversity ($\pi$) and Tajima's D (window size 10 Kbp, step size 2 Kbp) for spring and fall spawners, in light and dark blue lines, respectively; and estimate of recombination rate (rho/Kbp) every 100 Kbp (Pettersson et al., 2019). Zoom-in plots to 5 loci within this region, (**C**) chr15:6,750,000-7,000,000, (**D**) chr15:7,650,000-7,850,000, (**E**) chr15:8,540,000-9,070,000, (**F**) chr15:9,200,000-9,350,000, (**G**) chr15:10,820,000-11,100,000. Each plot has four tracks: gene models, dAF, nucleotide diversity, and Tajima's D.

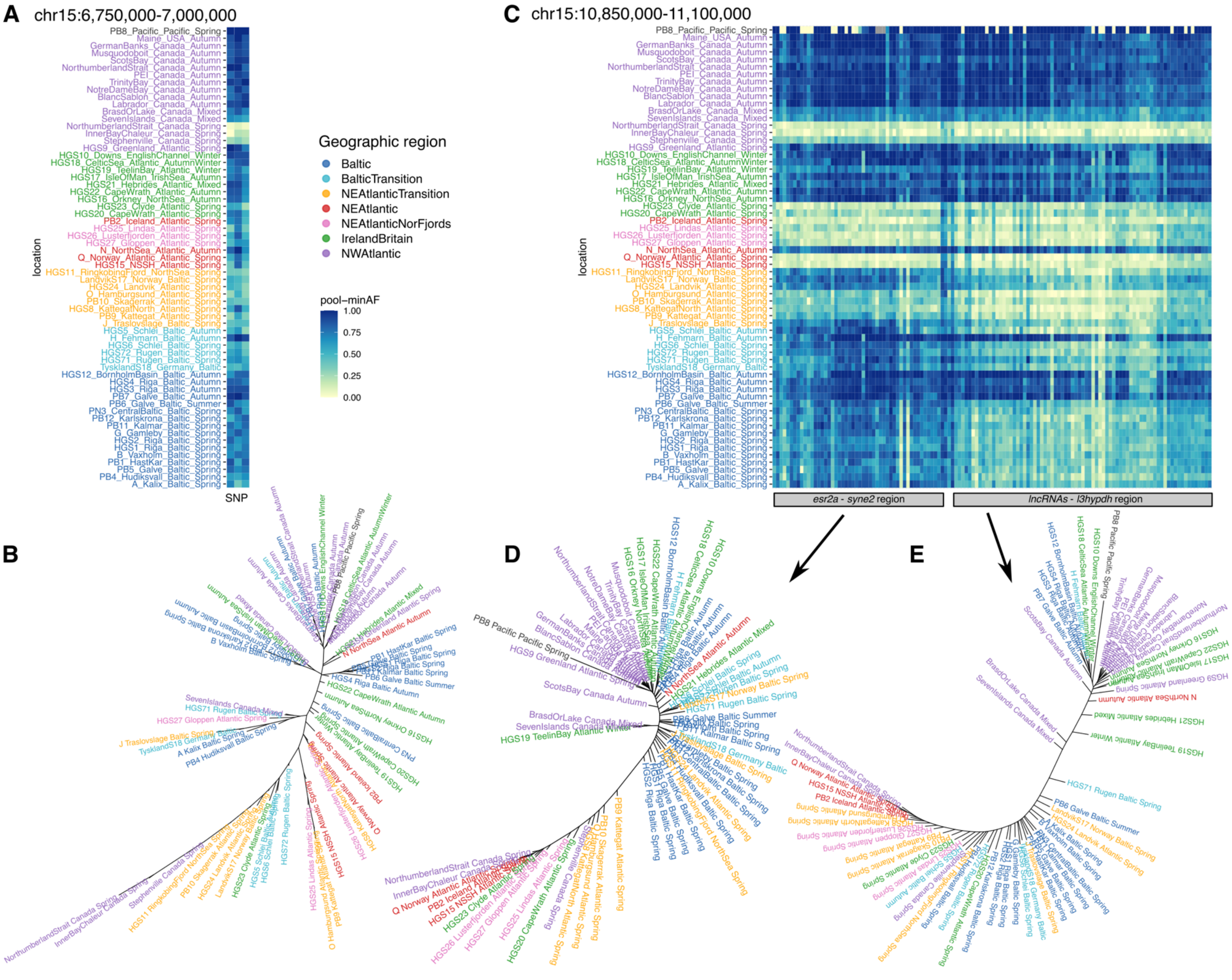
A
chr15:6,750,000-7,000,000
C
chr15:10,850,000-11,100,000
PB8_Pacific_Pacific_Spring
Maine_USA_Autumn
GermanBanks_Canada_Autumn
Musquodoboit_Canada_Autumn
ScotsBay_Canada_Autumn
NorthumberlandStrait_Canada_Autumn
PEI_Canada_Autumn
TrinityBay_Canada_Autumn
NotreDameBay_Canada_Autumn
BlancSablon_Canada_Autumn
Labrador_Canada_Autumn
BrasdOrLake_Canada_Mixed
SevenIslands_Canada_Mixed
NorthumberlandStrait_Canada_Spring
InnerBayChaleur_Canada_Spring
Stephenville_Canada_Spring
HGS9_Greenland_Atlantic_Spring
HGS10_Downs_EnglishChannel_Winter
HGS18_CelticSea_Atlantic_AutumnWinter
HGS19_TeelinBay_Atlantic_Winter
HGS17_IsleOfMan_IrishSea_Autumn
HGS21_Hebrides_Atlantic_Mixed
HGS22_CapeWrath_Atlantic_Autumn
HGS16_Orkney_NorthSea_Autumn
HGS23_Clyde_Atlantic_Spring
HGS20_CapeWrath_Atlantic_Spring
PB2_Iceland_Atlantic_Spring
HGS25_Lindas_Atlantic_Spring
HGS26_Lusterfjorden_Atlantic_Spring
HGS27_Gloppen_Atlantic_Spring
N_NorthSea_Atlantic_Autumn
Q_Norway_Atlantic_Atlantic_Spring
HGS15_NSSH_Atlantic_Spring
HGS11_RingkobingFjord_NorthSea_Spring
LandvikS17_Norway_Baltic_Spring
HGS24_Landvik_Atlantic_Spring
O_Hamburgsund_Atlantic_Spring
PB10_Skagerrak_Atlantic_Spring
HGS8_KattegatNorth_Atlantic_Spring
PB9_Kattegat_Atlantic_Spring
J_Traslovslage_Baltic_Spring
HGS5_Schlei_Baltic_Autumn
H_Fehmarn_Baltic_Autumn
HGS6_Schlei_Baltic_Spring
HGS72_Rugen_Baltic_Spring
HGS71_Rugen_Baltic_Spring
TysklandS18_Germany_Baltic
HGS12_BornholmBasin_Baltic_Autumn
HGS4_Riga_Baltic_Autumn
HGS3_Riga_Baltic_Autumn
PB7_Galve_Baltic_Autumn
PB6_Galve_Baltic_Summer
PN3_CentralBaltic_Baltic_Spring
PB12_Karlskrona_Baltic_Spring
PB11_Kalmar_Baltic_Spring
G_Gamleby_Baltic_Spring
HGS2_Riga_Baltic_Spring
HGS1_Riga_Baltic_Spring
B_Vaxholm_Baltic_Spring
PB1_HastKar_Baltic_Spring
PB5_Galve_Baltic_Spring
PB4_Hudiksvall_Baltic_Spring
A_Kalix_Baltic_Spring
location
SNP
Geographic region
Baltic
BalticTransition
NEAtlanticTransition
NEAtlantic
NEAtlanticNorFjords
IrelandBritain
NWAtlantic
pool−minAF
1.00
0.75
0.50
0.25
0.00
esr2a - syne2 region
lncRNAs - l3hypdh region
B
D
E

**Figure S10**. Comparison of the genetic patterns in two outlier regions in chr 15. The first region (**A-B**) harbors the *rab15-si* gene. (**A**) Pool-allele frequencies and (**B**) neighbor-joining tree based on the novel genetic variants (dAF > 0.55). The second region (**C-E**) comprises the *esr2a-syne2-lncRN-l3hypdh* genes. (**C**) Pool-allele frequencies of outlier SNPs for the entire region. neighbor-joining tree (**D**) for the novel genetic variants in the *esr2a-syne2* region, and in the (**E**) *lncRN-l3hypdh* region. Note that the sample from Greenland corresponds to summer spawners as indicated in the Supplementary file 1 of Han et al. (2020).

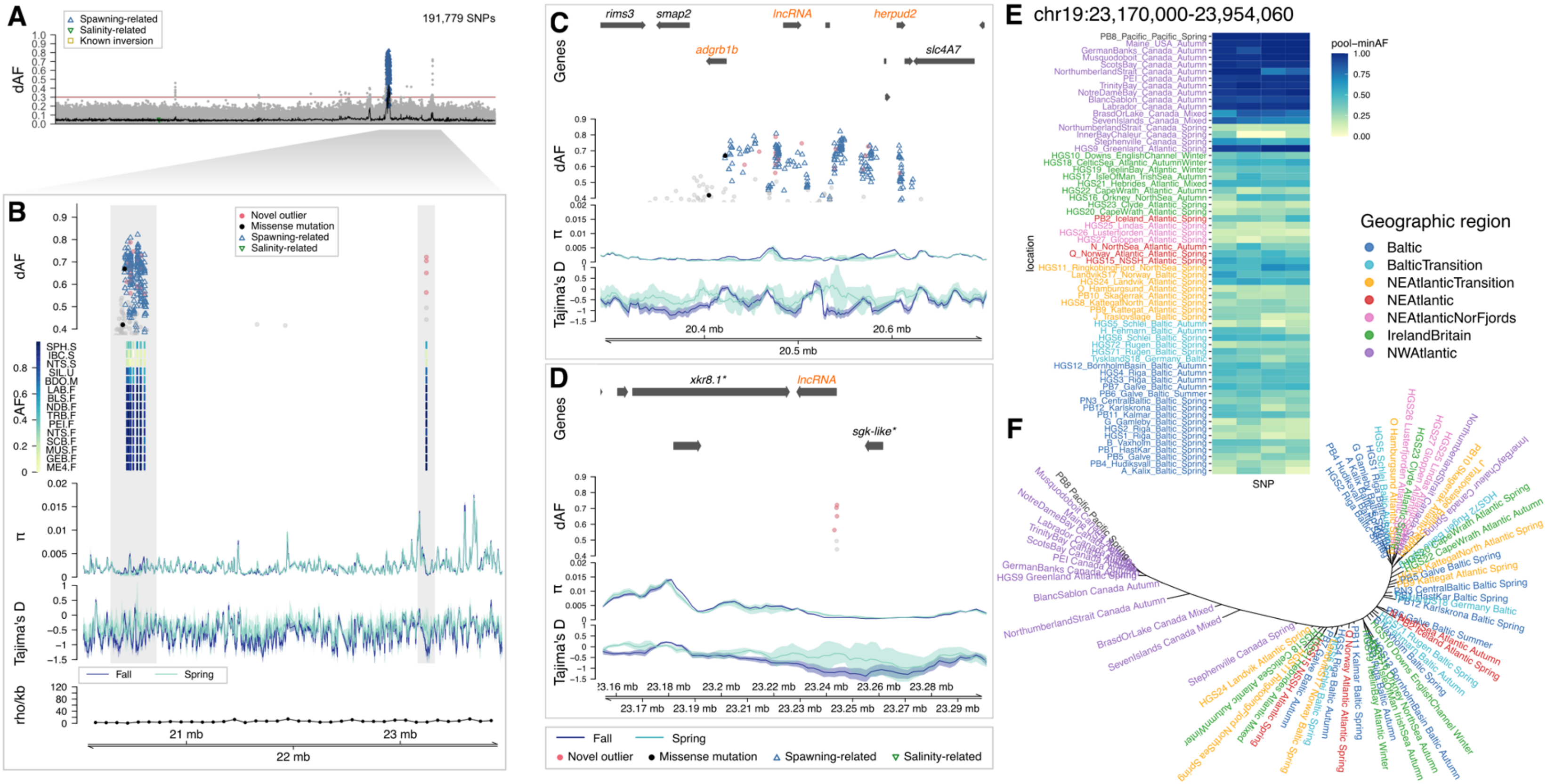


**Figure S11**. Selection signal on chr 19. (**A**) Genetic differentiation (dAF) along the chromosome. (**B**) Close-up to the target region. This plot consists of five tracks, from top to bottom: genetic differentiation between spring and fall spawners for SNPs with dAF ≥ 0.4. Novel outlier SNPs (dAF ≥ 0.55) are denoted as red filled circles, missense mutations as filled black circles, spawning-related SNPs as empty blue triangles, and other SNPs are gray circles; heatmap plot depicting the minor allele frequency per population (rows) for the novel outlier SNPs (columns); average nucleotide diversity ($\pi$) and Tajima's D (window size 10 Kbp, step size 2 Kbp) for spring and fall spawners, in light and dark blue lines, respectively; and estimate of recombination rate (rho/Kbp) every 100 Kbp (Pettersson et al., 2019). Zoom-in plots to 5 loci within this region, (**C**) 19:20,290,000-20,700,000, and (**D**) 19:23,155,000-23,300,000. Each plot has four tracks: gene models, dAF, nucleotide diversity, and Tajima's D. (**E**) Pool-allele frequencies of diagnostic SNPs (dAF >= 0.55) (**F**) neighbor-joining tree based on these genetic variants. Note that the sample from Greenland corresponds to summer spawners as indicated in the Supplementary file 1 of Han et al. (2020).

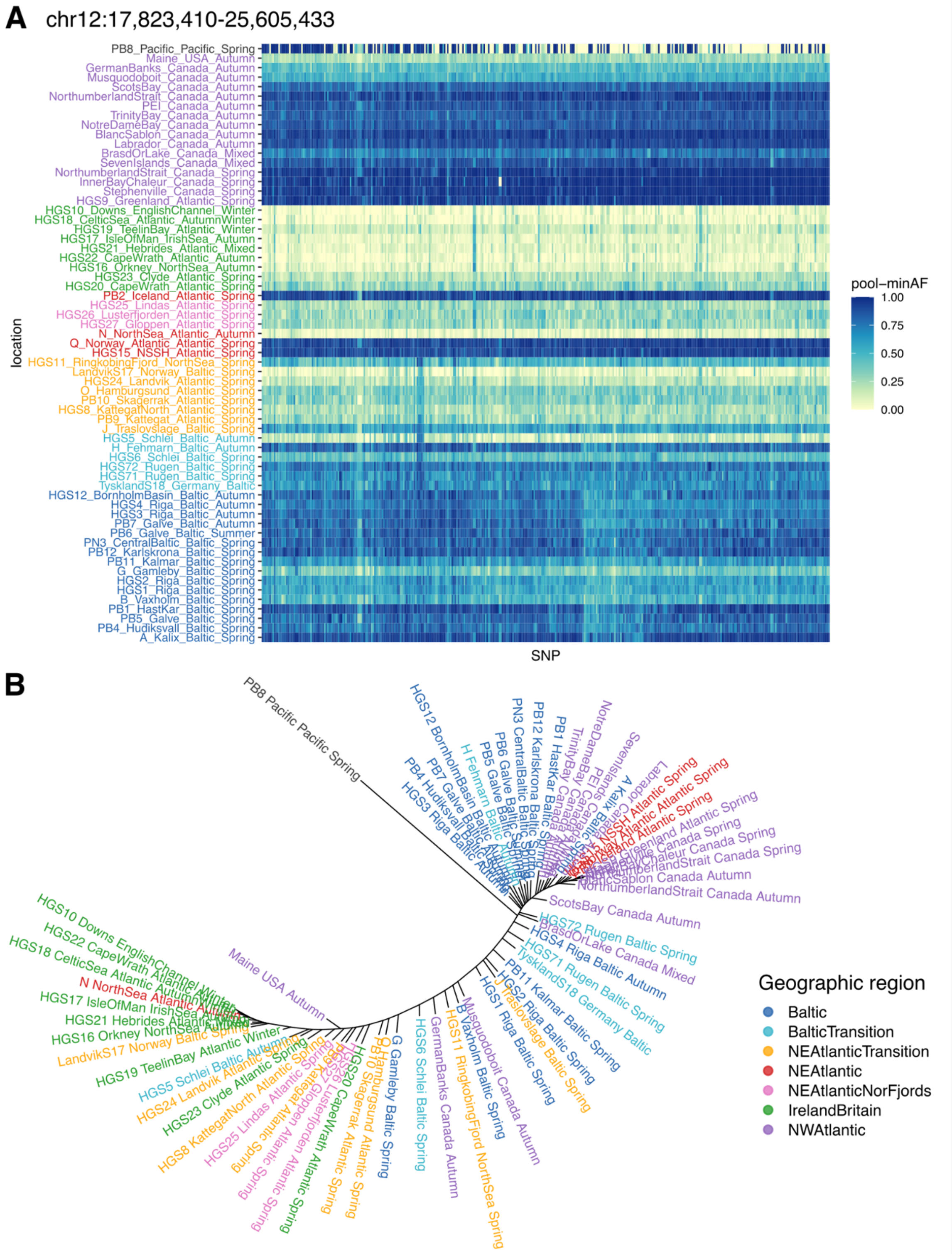


**Figure S12**. Selection signal on chr 12 corresponding to a known chromosomal inversion. (**A**) Pool-allele frequencies and (**B**) neighbor-joining tree based on diagnostic variants (dAF >= 0.45).

Note that the sample from Greenland corresponds to summer spawners as indicated in the Supplementary file 1 of Han et al. (2020).

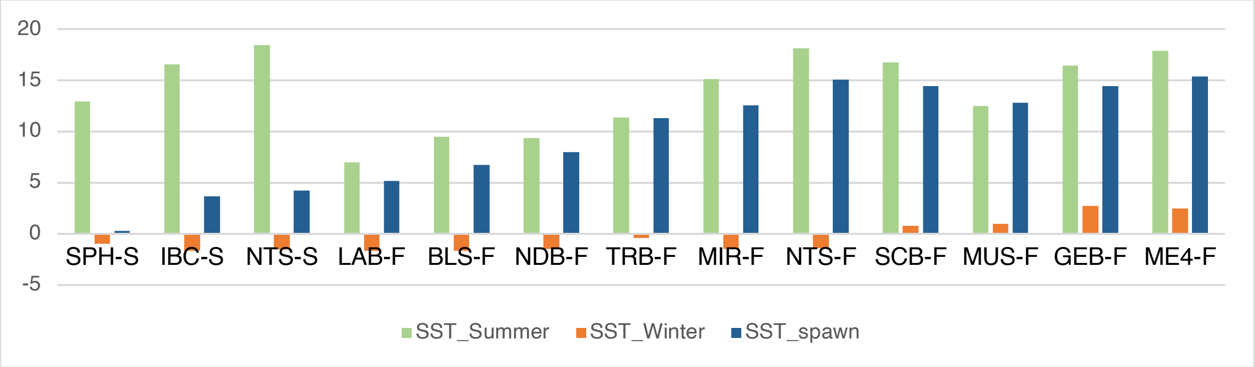


**Figure S13.** Average sea water temperature (°C) per location. Green bars indicate values for the summer months ($SST_{Summer}$), orange bars show values for the winter months ($SST_{Winter}$), and blue bars are for the spawning month ($SST_{spawn}$).

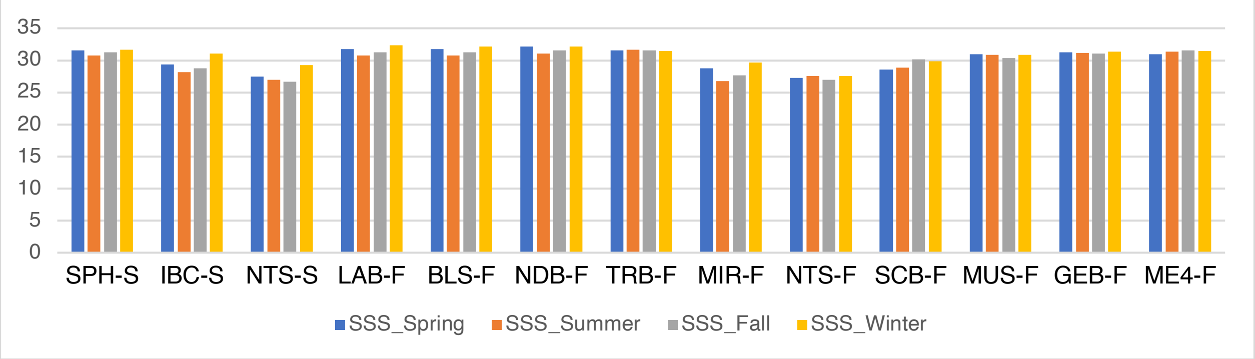


**Figure S14.** Average sea water salinity (PSU) per location. Blue bars indicate values for the summer months ($SST_{Summer}$), orange bars show values for the winter months ($SST_{Winter}$), and blue bars are for the spawning month ($SST_{spawn}$).

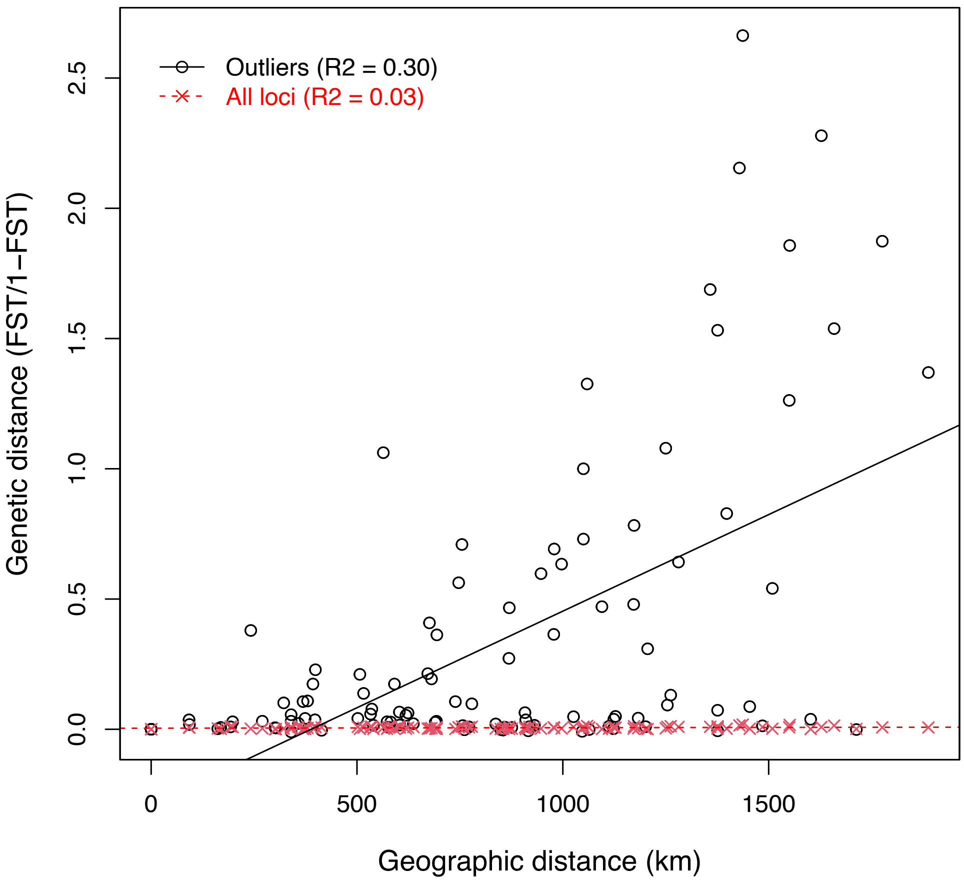


**Figure S15.** Examination of an Isolation-by-distance pattern for all SNPs (red x symbols) and 434 outliers SNPs (black empty circles, dAF >= 0.45) within the inversion on chr 12 showing a latitudinal pattern that differentiates northern and southern populations respect to a biogeographic transition zone in the Scotian shelf.
